# Shallow neural networks trained to detect collisions recover features of visual loom-selective neurons

**DOI:** 10.1101/2021.07.07.451307

**Authors:** Baohua Zhou, Zifan Li, Sunnie S. Y. Kim, John Lafferty, Damon A. Clark

**Author notes:** **For correspondence:** (DAC); (JDL). Department of Computer Science, Princeton University, Princeton, NJ, United States.

## Abstract

Animals have evolved sophisticated visual circuits to solve a vital inference problem: detecting whether or not a visual signal corresponds to an object on a collision course. Such events are detected by specific circuits sensitive to visual looming, or objects increasing in size. Various computational models have been developed for these circuits, but how the collision-detection inference problem itself shapes the computational structures of these circuits remains unknown. Here, inspired by the distinctive structures of LPLC2 neurons in the visual system of *Drosophila*, we build an anatomically-constrained shallow neural network model and train it to identify visual signals that correspond to impending collisions. Surprisingly, the optimization arrives at two distinct, opposing solutions, only one of which matches the actual dendritic weighting of LPLC2 neurons. The LPLC2-like solutions are favored when a population of units is trained on the task, but not when units are trained in isolation. The trained model reproduces experimentally observed LPLC2 neuron responses for many stimuli, and reproduces canonical tuning of loom sensitive neurons, even though the models are never trained on neural data. These results show that LPLC2 neuron properties and tuning are predicted by optimizing an anatomically-constrained neural network to detect impending collisions.

## Introduction

For animals living in dynamic visual environments, it is important to detect the approach of predators or other dangerous objects. Many species, from insects to humans, rely on a range of visual cues to identify approaching, or looming, objects [Regan and Beverley, 1978, Sun and Frost, 1998, Gabbiani et al., 1999, Card and Dickinson, 2008, Münch et al., 2009, Temizer et al., 2015]. Among other cues, looming objects create characteristic visual flow fields. When an object is on a ballistic collision course with an animal, its edges will appear to the observer to expand radially outward, gradually occupying a larger and larger portion of the visual field (*Figure 3, Video 1*). An object heading towards the animal, but which will not collide with it, also expands to occupy an increasing portion of the visual field, but its edges do not expand radially outwards with respect to the observer. Instead, they expand with respect to the object’s center so that opposite edges are perceived to be moving in the same direction (*Figure 3, Video 2*). A collision detector must distinguish between these two cases, while also avoiding predicting collisions in response to a myriad of other visual flow fields created by the animal’s own motion (*Figure 3, Video 4*). Thus, loom detection can be framed as a visual inference problem.

Many sighted animals solve this inference problem with high precision, thanks to robust loom-selective neural circuits evolved over hundreds of millions of years. The neuronal mechanisms for response to looming stimuli have been studied in a wide range of vertebrates, from cats and mice to zebrafish, as well as in humans [King et al., 1992, Hervais-Adelman et al., 2015, Ball and Tronick, 1971, Liu et al., 2011, Salay et al., 2018, Liu et al., 2011, Shang et al., 2015, Wu et al., 2005, Temizer et al., 2015, Dunn et al., 2016, Bhattacharyya et al., 2017]. In invertebrates, detailed anatomical, neurophysiological, behavioral and modeling studies have investigated loom detection, especially for locusts and flies [Oliva and Tomsic, 2014, Sato and Yamawaki, 2014, Santer et al., 2005, Rind and Bramwell, 1996, Card and Dickinson, 2008, De Vries and Clandinin, 2012, Muijres et al., 2014, Klapoetke et al., 2017, Von Reyn et al., 2017, Ache et al., 2019]. An influential mathematical model of loom detection was derived by studying the responses of the giant descending neurons of locusts, which established a relationship between the timing of the neurons’ peak responses and an angular size threshold for the looming object [Gabbiani et al., 1999]. Similar models have been applied to analyze neuronal responses to looming signals in flies, where genetic tools make it possible to precisely dissect neural circuits, revealing various neuron types that are sensitive to looming signals [Von Reyn et al., 2017, Ache et al., 2019, Morimoto et al., 2020]. However, these computational studies did not directly investigate the relationship between the structure of the loom-sensitive neural circuits and the inference problem they appear to solve. Here, we asked whether we can achieve the properties associated with neural loom detection simply by optimizing shallow neural networks for collision detection.

The starting point for our computational model of loom detection is the known neuroanatomy of the visual system of the fly. In particular, the loom-sensitive neuron LPLC2 (lobula plate/lobula columnar, type 2) [Wu et al., 2016] has been studied in detail. These neurons tile visual space, sending their axons to a descending neuron called the giant fiber (GF), which triggers the fly’s jumping and take-off behaviors [Tanouye and Wyman, 1980, Card and Dickinson, 2008, Von Reyn et al., 2017, Ache et al., 2019]. Each LPLC2 neuron has four dendritic branches that receive inputs from the four layers of the lobula plate (LP) (*Figure 1A*) [Maisak et al., 2013, Klapoetke et al., 2017]. The retinotopic LP layers host the axon terminals of motion detection neurons, and each layer uniquely receives motion information in one of the four cardinal directions [Maisak et al., 2013]. Moreover, the physical extensions of the LPLC2 dendrites align with the preferred motion directions in the corresponding LP layers (*Figure 1B*) [Klapoetke et al., 2017]. These dendrites form an outward radial structure, which matches the moving edges of a looming object that expands in the visual field (*Figure 1*C). Common stimuli such as the wide-field motion generated by movement of the insect only match part of the radial structure, and strong inhibition for inward-directed motion suppresses responses to such stimuli. Thus, the structure of the LPLC2 dendrites favors responses to objects with edges moving radially outwards, corresponding to motion toward center of the receptive field.

**Figure 1.**
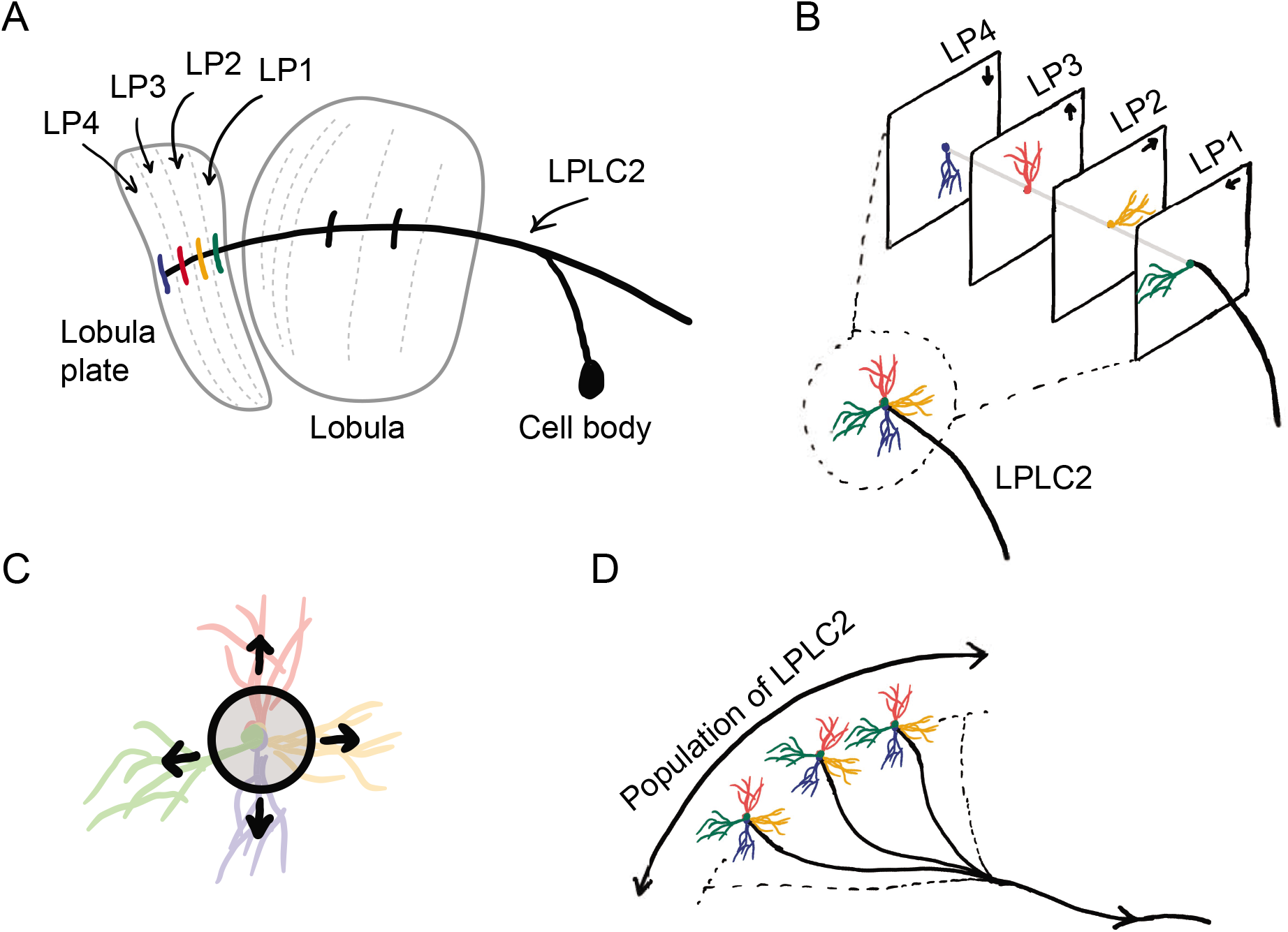
Sketches of the anatomy of LPLC2 neurons [Klapoetke et al., 2017]. (A) An LPLC2 neuron has dendrites in lobula and the four layers of the lobula plate (LP): LP1, LP2, LP3 and LP4. (B) Schematic of the four branches of the LPLC2 dendrites in the four layers of the LP. The arrows indicate the preferred direction of motion sensing neurons with axons in each LP layer [Maisak et al., 2013]. (C) The outward dendritic structure of an LPLC2 neuron is selective for the outwardly expanding edges of a looming object (black circle). (D) The axons of a population of more than 200 LPLC2 neurons converge to the GF, a descending neuron, to contribute to signaling for escaping behaviors [Ache et al., 2019]

The focus of this paper is to investigate how loom detection in LPLC2 can be seen as the solution to a computational inference problem. Can the structure of the LPLC2 neurons be explained in terms of optimization—carried out during the course of evolution—for the task of predicting which trajectories will result in collisions? How does coordination among the population of more than 200 LPLC2 neurons tiling a fly’s visual system affect this optimization? To answer these questions, we built a simple anatomically-constrained neural network model, which receives motion signals in the four cardinal directions. We trained the model to detect visual objects on a collision course with the observer using artificial stimuli. Surprisingly, optimization finds two distinct types of solutions, with one resembling the LPLC2 neurons and the other having a very different configuration. We analyzed how each of these solutions detects looming events and where they show distinct individual and population behaviors. When the number of units tiling visual space is increased, the solutions that resemble the actual LPLC2 neurons become favored. When tested on visual stimuli not in the training data, the optimized solutions exhibit response curves that are similar to those of actual LPLC2 neurons as measured by Klapoetke et al. [2017]. Importantly, the optimized model reproduces the canonical linear relationship between the timing of the peak responses and the size-to-speed ratio [Gabbiani et al., 1999]. Although only receiving motion signals, the model shows characteristics of an angular size encoder, which is consistent with many biological loom detectors [Gabbiani et al., 1999, Von Reyn et al., 2017, Ache et al., 2019]. Our results show that optimizing a neural network to detect looming events gives rise to the properties and tuning of LPLC2 neurons.

## Results

### A set of artificial visual stimuli is designed for training models

Our goal is to compare computational models trained to perform loom-detection with the biological computations in LPLC2 neurons. We first created a set of stimuli to act as training data for the inference task (Methods and Materials). We considered the following four types of motion stimuli: loom-and-hit (abbreviated as hit), loom-and-miss (miss), retreat, and rotation (*Figure 2*). The hit stimuli consist of a sphere that moves ballistically towards the origin on a collision course (*Video 1*). The miss stimuli consist of a sphere that moves ballistically towards the origin but misses it (*Video 2*). The retreat stimuli consist of a sphere moving ballistically away from the origin (*Video 3*). The rotation stimuli consist of objects rotating about an axis going through the origin (*Video 4*). All stimuli were designed to be isotropic; the first three stimuli could have any orientation in space, while the rotation could be about any axis through the origin. All trajectories were simulated in the frame of reference of the fly at the origin, with distances measured with respect to the origin. For simplicity, the fly is assumed to be a point particle with no volume (Red dots in *Figure 2* and the apexes of the cones in *Figure 3*). For hit, miss, and retreat stimuli, the spherical object has unit radius, and for the case of rotation, there were 100 objects of various radii scattered isotropically around the fly (*Figure 3*).

**Figure 2.**
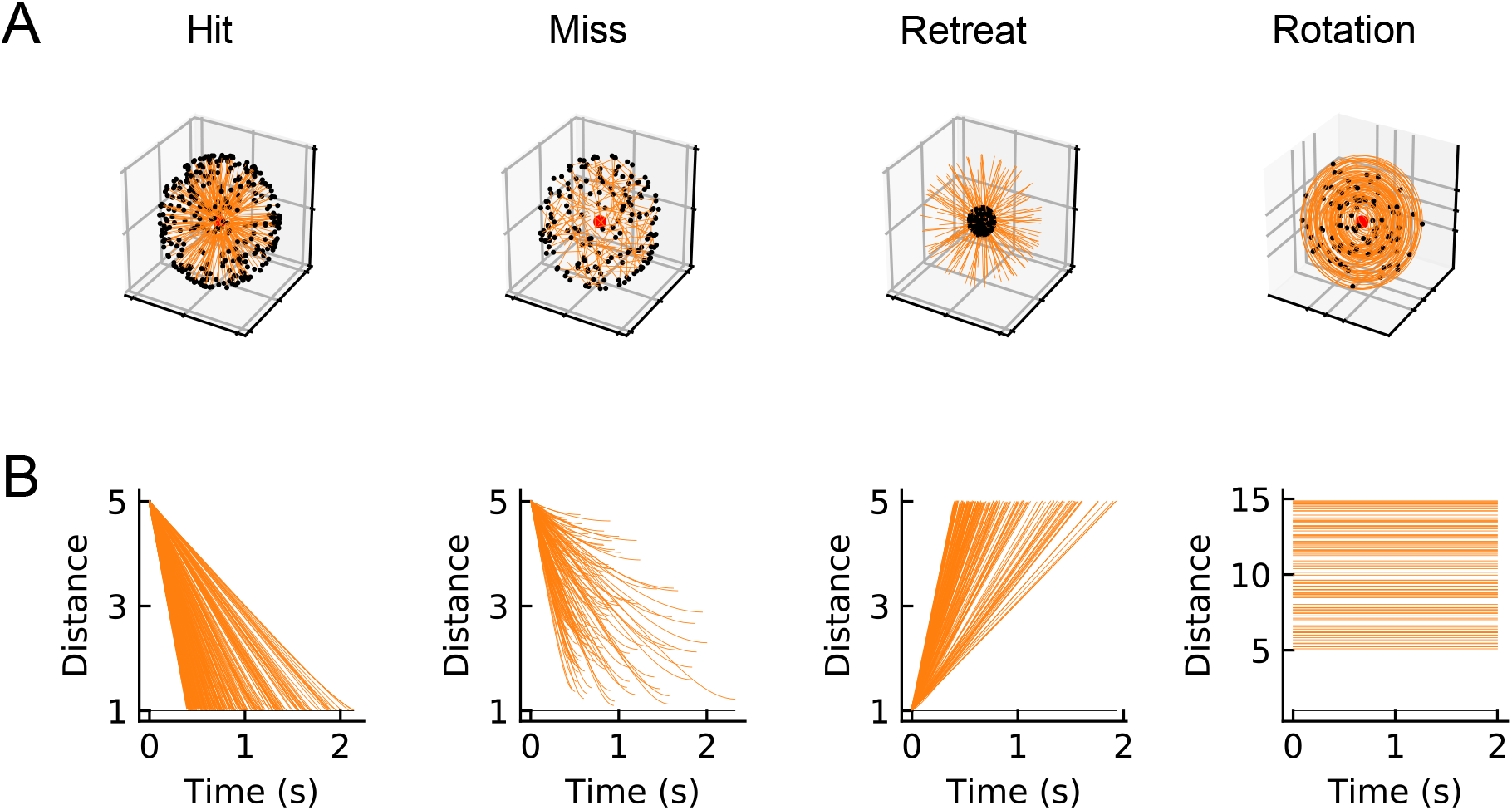
Four types of synthetic stimuli (Methods and Materials). (A) Orange lines represent trajectories of the stimuli. The black dots represent the starting points of the trajectories. For hit, miss, and retreat cases, multiple trajectories are shown. For rotation, only one trajectory is shown. (B) Distances of the objects to the fly eye as a function of time. Among misses, only the approaching portion of the trajectory was used. The horizontal black lines indicate the distance of 1.

**Figure 3.**
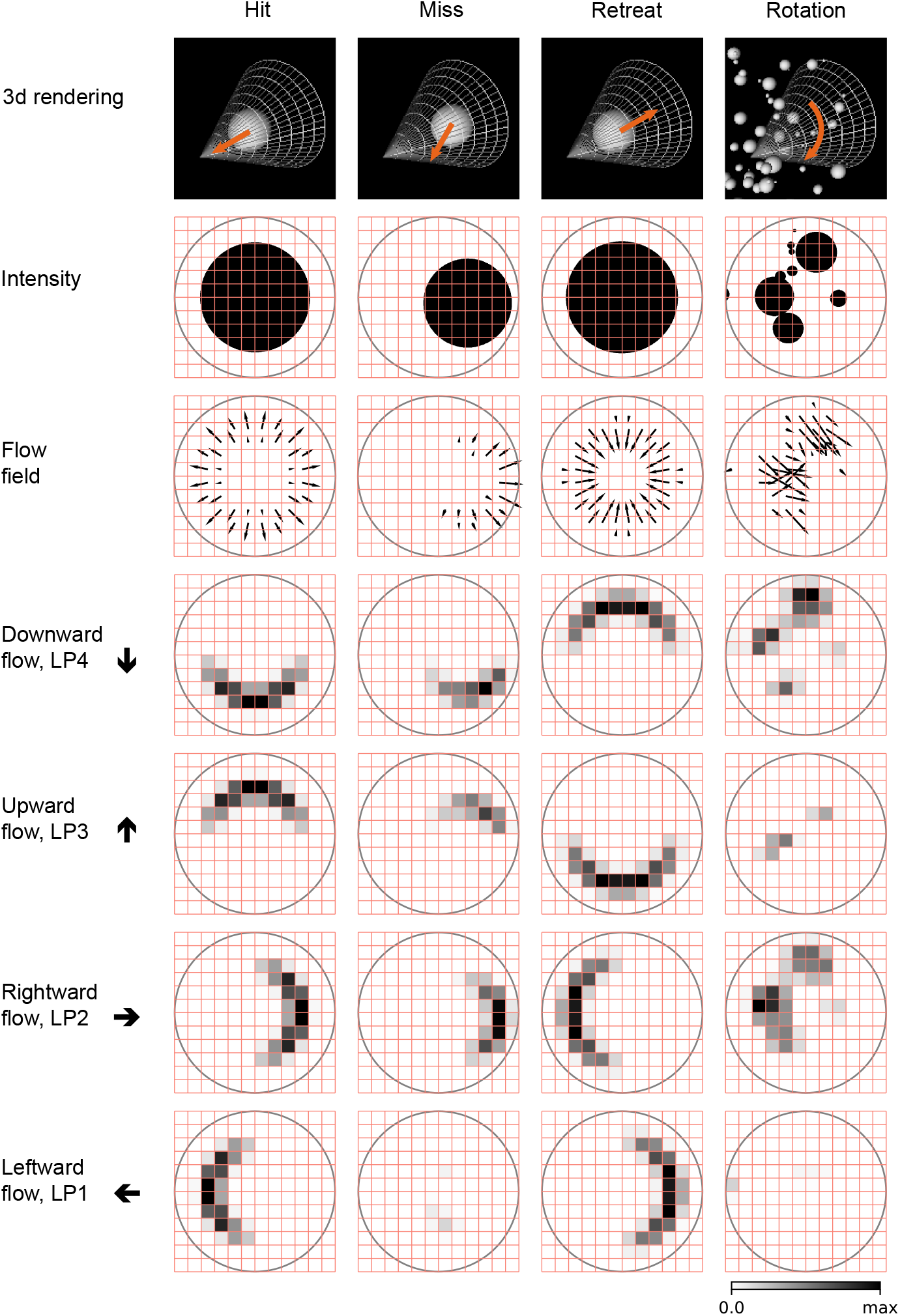
Snapshots of optical flows and flow fields calculated by a Hassenstein Reichardt correlator (HRC) model (Methods and Materials) for the 4 types of stimuli (*Figure 2*). First row: 3d rendering of the spherical objects and the LPLC2 receptive field (represented by a cone) at a specific time in the trajectory. The orange arrows indicate the motion direction of each object. Second row: 2d projections of the objects (black shading) within the LPLC2 receptive field (the grey circle). Third row: the thin black arrows indicate flow fields generated by the edges of the moving objects. Forth to seventh rows: decomposition of the flow fields in the four cardinal directions with respect to the LPLC2 neuron under consideration: downward, upward, rightward, and leftward, as indicated by the thick black arrows. These act as models of the motion signal fields in each layer of the lobula plate. **Figure 3–Figure supplement 1.** Tuning curve of HRC motion estimator and distributions of the estimated flow fields.

**Video 1.** Movie for a hit stimulus (single unit). Top left panel: 3d rendering as in the top row of *Figure 3*; bottom left panel: optical signal as in the second row of *Figure 3*; top right panel: flow fields in the horizontal direction as in rows 7 and 8 of *Figure 3*; bottom right panel: flow fields in the vertical direction as in rows 5 and 6 of *Figure 3*. Since we combined left (down) and right (up) flow fields in one panel, we used blue and red colors to indicate left (down) and right (up) directions, respectively. The movie has been slowed down by a factor of 5. All the movies shown in this paper can be found here: https://github.com/ClarkLabCode/LoomDetectionANN/tree/main/results/movies_exp.

**Video 2.** Movie for a miss stimulus (single unit). The same arrangement as *Video 1*.

**Video 3.** Movie for a retreat stimulus (single unit). The same arrangement as *Video 1*.

**Video 4.** Movie for a rotation stimulus (single unit). The same arrangement as *Video 1*.

### An anatomically-constrained mathematical model

We designed and trained a simple, anatomically-constrained neural network (*Figure 4* to infer whether or not a moving object will collide with the fly. The features of this network were designed to mirror anatomical features of the fly’s LPLC2 neurons (*Figure 1*). Model units receive input from a 60 degree diameter cone of visual space, represented by white cones and grey circles in *Figure 3*, mirroring the receptive field size that has been measured for LPLC2 [Klapoetke et al., 2017]. The four stimulus sets were projected into this receptive field for training and evaluating the model. The inputs to the model are local directional signals computed in the four cardinal directions at each point of the visual space: downward, upward, rightward, and leftward (*Figure 3*). These represent the motion signals from T4 and T5 neurons in the four layers of the lobula plate [Maisak et al., 2013]. They are computed as the non-negative components of a Hassenstein-Reichardt correlator model [Hassenstein and Reichardt, 1956] in both horizontal and vertical directions, which acts on the intensities of the projected stimuli (Methods and Materials). The motion signals are computed with a spacing of 5 degrees, roughly matching the spacing of the ommatidia and processing columns in the fly eye [Stavenga, 2003].

**Figure 4.**
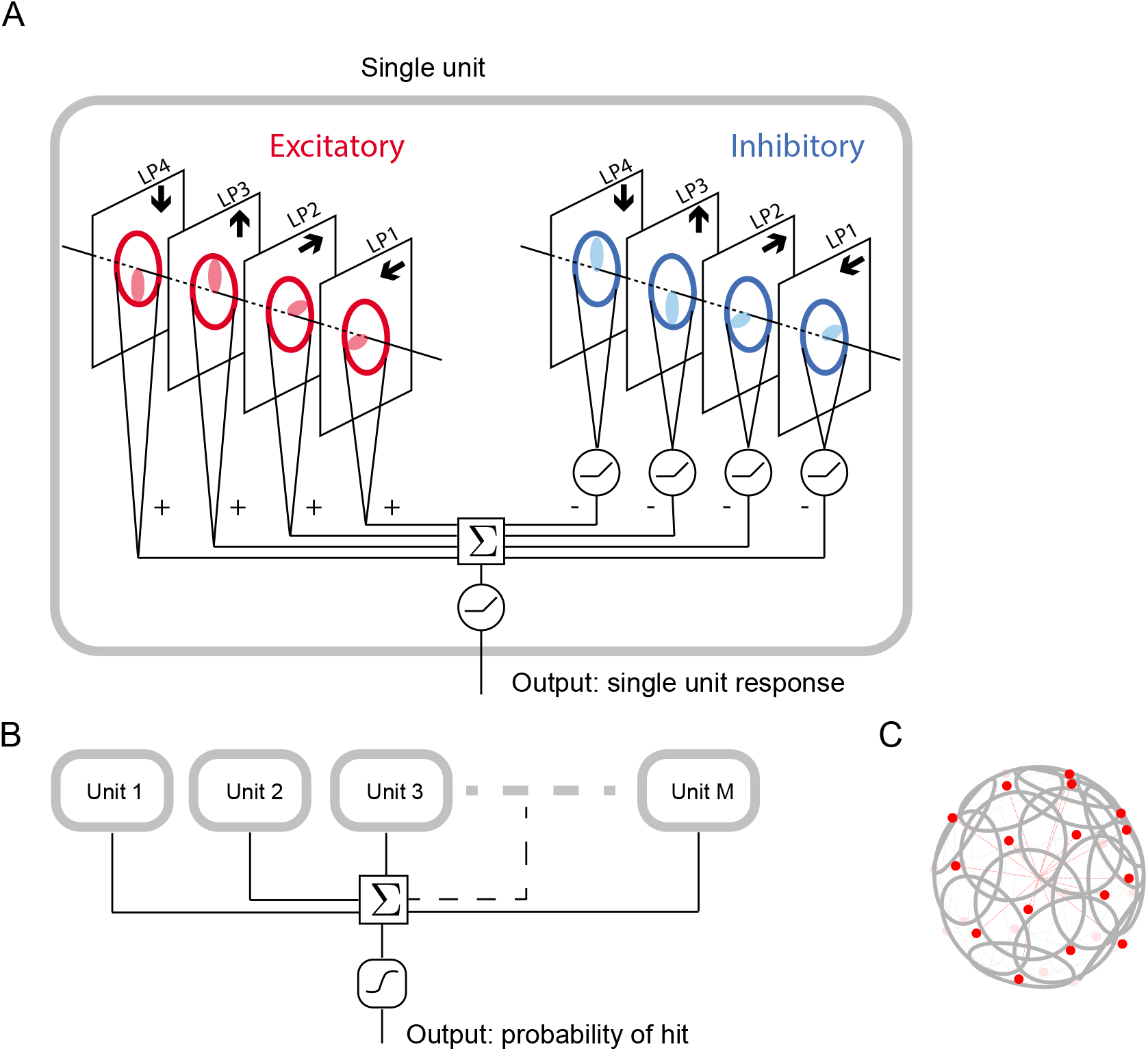
Schematic of the model (Methods and Materials). (A) Single unit. There are two sets of nonnegative filters: excitatory (red) and inhibitory (blue). Each set of filters has four branches, and each branch receives a field of motion signals (forth to seventh rows in *Figure 3*) from the corresponding layer of the model LP. The weighted signals from the excitatory branches and the inhibitory branches (rectified) are pooled together to go through a rectifier to produce an output, which is the response of a single unit. (B) The outputs from *M* units are summed and fed into a sigmoid function to estimate the probability of hit. (C) The *M* units have their orientations almost evenly distributed in angular space. Red dots represent the centers of the receptive fields and the grey lines represent the boundaries of the receptive fields on unit sphere. The red lines are drawn from the origin to the center of each receptive field. **Figure 4–Figure supplement 1.** Coordinate system for model and stimuli.

Each model unit can weight the motion signals from the four layers using linear spatial filters. There are two sets of non-negative filters, the excitatory filters and the inhibitory filters; these are shown in red and blue, respectively (*Figure 4*A). Each set of filters has four components, or branches, integrating motion signals from the four cardinal directions, respectively. These spatial filters represent excitatory inputs to LPLC2 directly from T4 and T5 in the LP, and inhibitory inputs mediated by local interneurons [Klapoetke et al., 2017, Mauss et al., 2015]. All eight filters act on the 60 degree receptive field of an unit. A 90-degree rotational symmetry is imposed on the filters, so that the filters in each layer are identical. Moreover, each filter is symmetric about the axis of motion (Methods and Materials). No further assumptions were made about the structures of the filters.

The model incorporates a fundamental difference between the excitatory and inhibitory branches: while the integrated signals from each excitatory branch are sent directly to the downstream computations, the integrated signals from each inhibitory branch are rectified before being sent downstream. This difference reflects anatomical constraints of the inputs to an actual LPLC2 neuron, where the excitatory inputs are direct connections with LPLC2 while the inhibitory inputs are mediated by inhibitory interneurons (LPi) between LP layers [Mauss et al., 2015, Klapoetke et al., 2017]. The outputs of the eight branches are summed and rectified to generate the output of a single model unit in response to a given stimulus (*Figure 4*A.)

In the fly brain, a population of LPLC2 neurons converges onto the GF (*Figure 1*D). Accordingly, in our model there are *M* replicates of model units, with orientations that are spread uniformly over the 4*π* steradians of the unit sphere (*Figure 4*C, Methods and Materials). In this way, the receptive fields of the *M* units roughly tile the whole angular space, with or without overlap, depending on the value of *M*. The sum of the responses of the *M* model units is fed into a sigmoid function to generate the predicted probability of collision for a given trajectory (Methods and Materials).

### Optimization finds two distinct solutions to the loom-inference problem

The objective of this study is to investigate how the binary classification task shapes the excitatory and inhibitory filters, and how the number of units *M* affects the results. We begin with the simplest model, which possess only a single unit, i.e., *M* =1. After training with 200 random initializations of the filters, we find that the converged solutions fall into three broad categories (*Figure 5*A, B). One set of solutions is largely unstructured, with almost all the elements in the filters equal to zero (labeled in black); we will ignore these for the rest of the analysis. The two structured solutions are interesting because, surprisingly, they have spatial structures that are roughly opposite from one another (magenta and green). Based on the configurations of the excitatory filters (Methods and Materials), we call one solution type *outward filters* (magenta), and the other type *inward filters* (green) (*Figure 5*C and *Figure 5–Figure Supplement 1*A). In this single-unit model, the inward solutions perform better than the outward solutions on the discrimination task (*Figure 5*D and *Figure 5–Figure Supplement 1*B).

**Figure 5.**
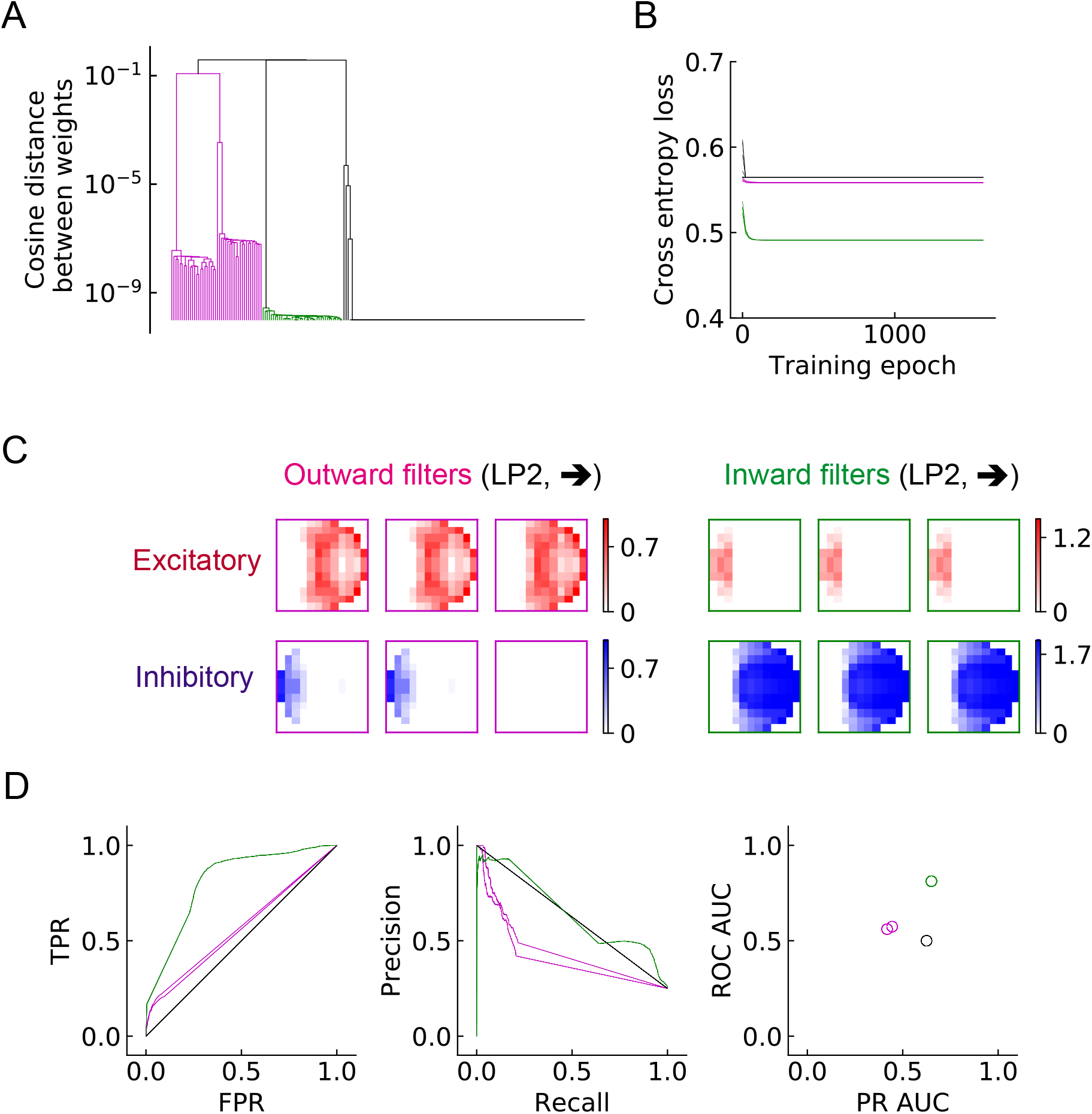
Two distinct types of solutions appear from training a single unit on the binary classification task. (A) Clustering of the trained filters/weights shown as a dendrogram (Methods and Materials). Different colors indicate different clusters, which are preserved for the rest of the paper (see (C)) (B) The trajectories of the loss functions during training. (C) The two distinct types of solutions are represented by two types of filters that have roughly opposing structures: an outward solution (magenta) and an inward solution (green). The excitatory filter weights are shown in red, and the inhibitory filters are shown in blue. (D) Performance of the two solution types (Methods and Materials). TPR: true positive rate; FPR: false positive rate; ROC: receiver operating characteristic; PR: precision recall; AUC: area under the curve. **Figure 5–Figure supplement 1.** More examples of the trained filters for the two types of solutions and the inferred probability of hit for the four types of training stimuli.

As the number of units *M* increases, the population of units covers a larger angular space, and when *M* is large enough (*M* ≥ 16), the receptive fields of the units begin to overlap with each other (*Figure 6*A). In the fly visual system there are over 200 LPLC2 neurons across both eyes [Ache et al., 2019], which corresponds to a very dense distribution of the units. This is illustrated by the third row in *Figure 6*A) where *M* = 256. When *M* is large, approaching objects from any direction are detectable and in fact, such object signals can be detected simultaneously by many neighboring units.

**Figure 6.**
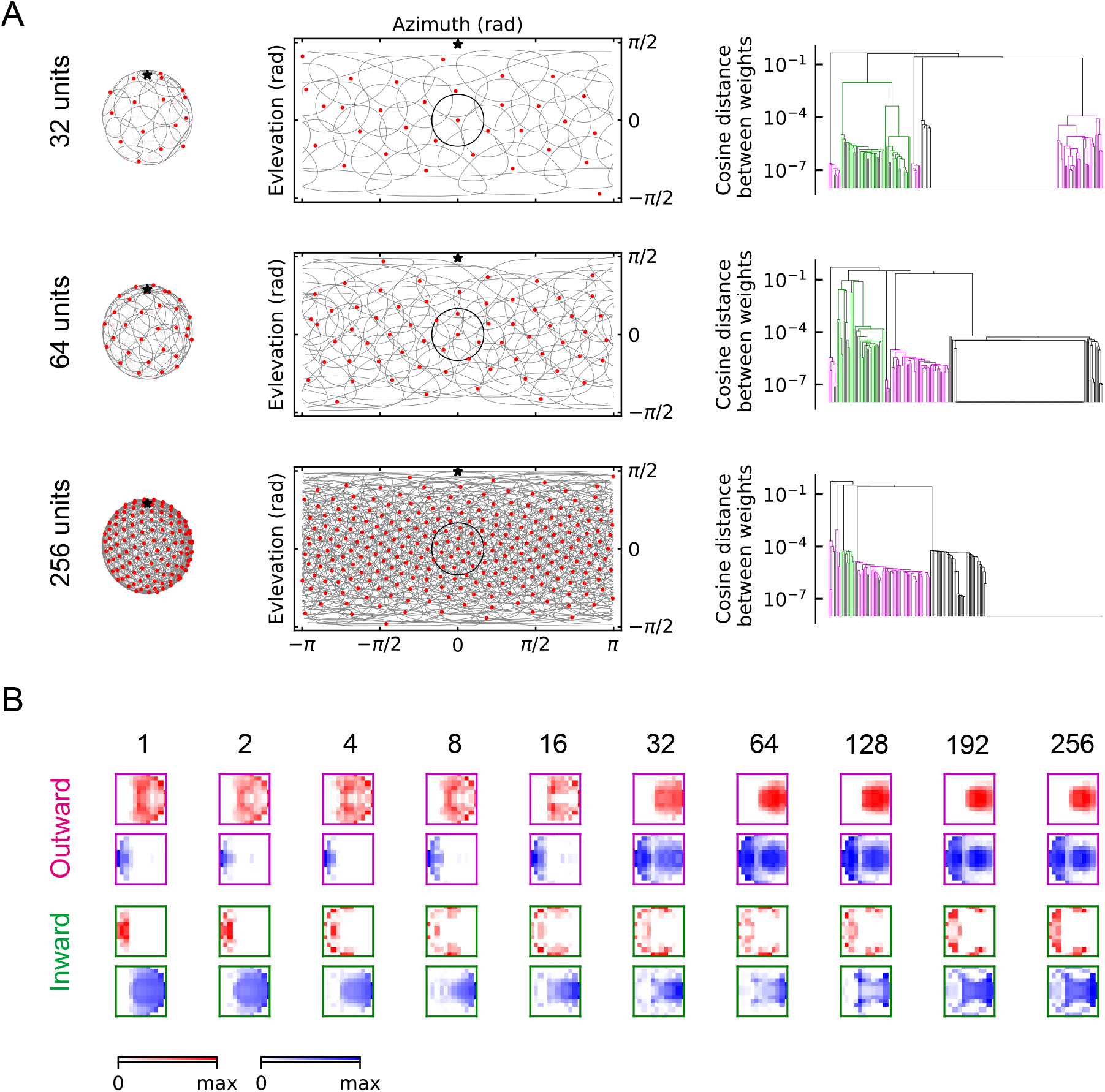
The outward and inward solutions also arise for models with multiple units. (A) Left column: angular distribution of the units, where red dots are centers of the receptive fields, the grey circles are the boundaries of the receptive fields, with one field highlighted in black, and the black star indicates the top of the fly head. Middle column: 2d map of the units with the same symbols as in the left column. Right column: clustering results shown as dendrogams with color codes as in *Figure 5*. (B) Examples of the trained excitatory and inhibitory filters for outward and inward solutions with different numbers of units. **Figure 6–Figure supplement 1.** Performance of the different solutions. **Figure 6–Figure supplement 2.** More examples of the outward filters. **Figure 6–Figure supplement 3.** More examples of the inward filters.

Interestingly, the two oppositely structured solutions persist, regardless of the value of *M* (*Figure 6, Figure 6–Figure Supplement 1, Figure 6–Figure Supplement 2, Figure 6–Figure Supplement 3*). In some outward solutions, structures on the right side of the inhibitory filters are similar to structures of the corresponding excitatory filters. This indicates a degree of redundancy, or non-identifiability in the model (*Figure 6–Figure Supplement 2, Figure 6–Figure Supplement 3*).

Units with outward-oriented filters are activated by motion radiating outwards from the center of the receptive field. Thus, these excitatory filters resemble the dendritic structures of the actual LPLC2 neurons observed in experiments, where for example, the rightward motion sensitive branch (LP2) occupies mainly the right side of the receptive field. In the outward solutions, the rightward motion-sensitive inhibitory filter mainly occupies the *left* side of the receptive field. This is also consistent with the properties of the lobula plate intrinsic (LPi) interneurons, which project inhibitory signals roughly retinotopically from one LP layer to an adjacent layer with opposite directional tuning [Mauss et al., 2015, Klapoetke et al., 2017].

The unexpected inward-oriented filters have the opposite structure. In the inward solutions, the rightward sensitive excitatory filter occupies the left side of the receptive field, and the inhibitory filter occupies the right side. Such weightings make the model selective for motion converging towards the receptive field center. At first glance, this is a puzzling structure for a loom detector, so we explored the response properties of the inward and outward solutions in more detail.

### Outward and inward filters are selective to signals in different ranges of angles

To understand the differences between the two types of solutions and why the inward filters can predict collisions, we investigated how they respond to hit stimuli from different incoming angles *θ* (*Figure 7*A). When there is no signal, the baseline activity of outward units is zero; however, the baseline activity of inward units is above zero (grey dashed lines in *Figure 7*B, C). This is because the trained intercepts are negative in the outward case, but positive in the inward case (Methods and Materials). Second, the outward filters respond strongly to stimuli near the center of the receptive field, but do not respond to stimuli having angles larger than ~ 30° (*Figure 7*B, C). In contrast, units with inward filters respond negatively to hit stimuli approaching toward the center and positively to stimuli approaching from the periphery of the receptive field, with angles between ~ 30° and ~ 90° (*Figure 7*B, C). This helps explain why the inward units can act as loom detectors: they are sensitive to hit stimuli originating in a larger solid angle. The hit signals are isotropic (*Figure 2*A), so the number of stimuli within angles ~ 30° and ~ 90° is much larger than the number of stimuli with angles below ~ 30° (*Figure 7*D). Thus, the inward solutions are sensitive to more hit cases than the outward solutions. One may visualize these responses as heat maps of the mean response of the models in terms of object distance to the fly and the incoming angle (*Figure 7*E). For the hit cases, the response patterns are consistent with the intuition about trajectory angles (*Figure 7*C). As expected, the inward solutions respond to the retreating signals with angles near ~ 180°, since the motion of edges in that case is radially inward.

**Figure 7.**
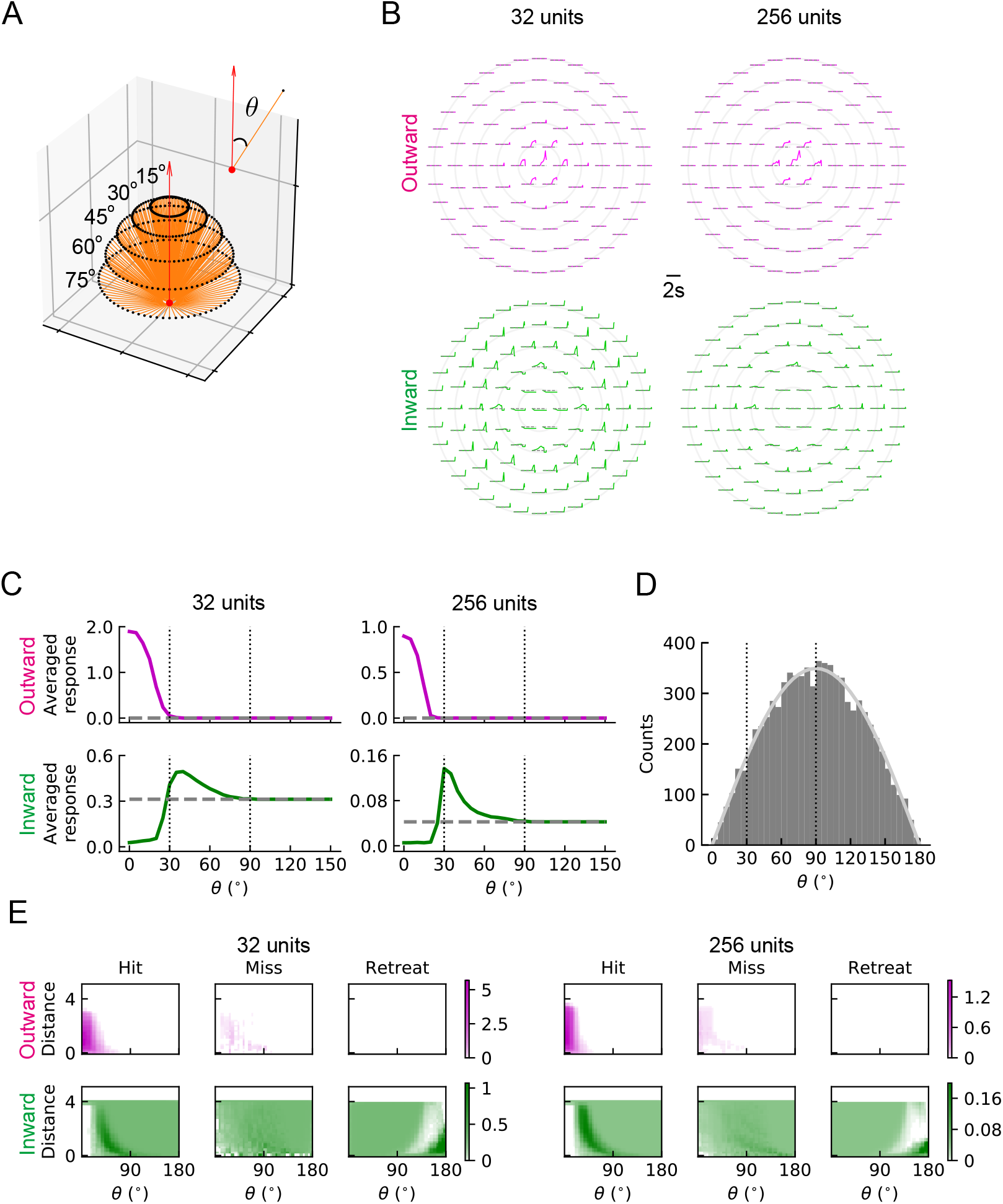
The outward and inward filters show distinct behaviors: single unit analysis. (A) Trajectories of hit stimuli with different incoming angles *θ*. Symbols are the same as in *Figure 2* except that the upward red arrow represents the orientation of one unit. The numbers with degree units indicate the specific values of the incoming angles. (B) Response patterns of a single unit with either outward (magenta) or inward (green) filters obtained from optimized solutions with 32 and 256 units, respectively. The grey dashed lines show the baseline activity of the unit when there is no stimulus. The solid grey concentric circles correspond to the values of the incoming angles in (A). The scale of the responses in the top left panel is four times the scale in the other three panels. (C) Temporally averaged responses against the incoming angle *θ*. Symbols and colors are the same as in (B). (D) Histogram of the incoming angles for the hit stimuli in *Figure 2*A. The grey curve represents a scaled sine function equal to the expected probability for isotropic stimuli. (E) Heatmaps of the response of a single unit against the incoming angle *θ* and the distance to the fly head, for both outward and inward filters obtained from optimized models with 32 and 256 units, respectively.

### Outward solutions have sparse codings and populations of units accurately predict hit probabilities

Individual units of the two solutions are very different from each other in their filter structure and response patterns to different stimuli. We decided to investigate how these differences manifest in the activities of populations of units, when units are trained to collectively predict the probability of hit. In populations of units, the outward and inward solutions exhibit very different response patterns for a given hit stimulus (*Figure 8*A, B, *Video 5* and *Video 6*). In particular, active outward units usually respond more strongly than inward units, but more inward units will be activated. This is consistent with the findings above, in which inward filter shapes responded to hits arriving from a wider distribution of angles.

**Figure 8.**
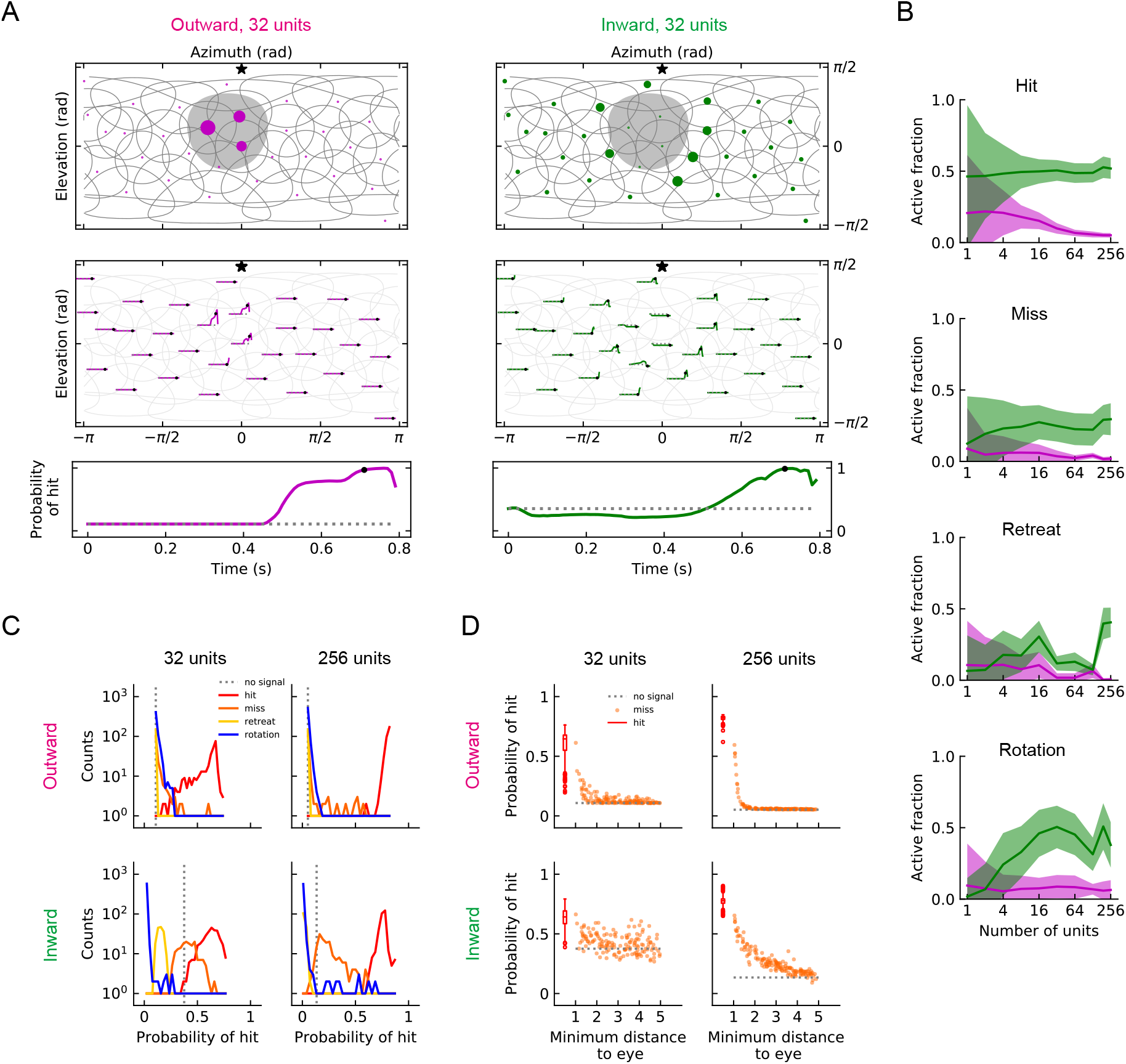
Population coding of stimuli. (A) Top row: a snapshot of the responses of outward units (magenta dots) for a hit stimulus (grey shade) (*Video 5* and *Video 6*). Symbols and colors are as in *Figure 6*A. Middle row: the whole trajectories of the responses for the same hit stimulus as in the top row. Bottom row: the entire trajectories of the probability of hit for the same hit stimulus as in the top row (Methods and Materials). Black dots in the middle and bottom rows indicate the time step of the snapshot in the top row. (B) Fractions of the units that are activated by different types of stimuli (hit, miss, retreat, rotation) as a function of the number of units *M* in the model. The lines represent the mean values averaged across samples, and the shaded areas show one standard deviation (Methods and Materials). (C) Histograms of the probability of hit inferred by models with 32 or 256 units for the four types of synthetic stimuli (Methods and Materials). (D) The inferred probability of hit as a function of the minimum distance of the object to the fly eye for the miss cases. The hit distribution is represented by a box plot (the center line in the box: the median; the upper and lower boundaries of the box: 25% and 75% percentiles; the upper and lower whiskers: the minimum and maximum; the circles: outliers). **Figure 8–Figure supplement 1.** Geometry of responses (as in Fig. *Figure 8*A), but for miss and retreat stimuli. **Figure 8–Figure supplement 2.** Sample individual unit response curves.

When a population of units encodes stimuli, at each time point, the sum of the activities of the units is used to infer the probability of hit. In our trained models, the outward and inward solutions give similar probabilities of hit (*Figure 8*A). Both types of solutions can give accurate inferences for the different stimuli (*Figure 8*C). In some cases, misses can be very similar to hits if the object passes near the origin. The models reflect this in their responses to near misses which have higher hit probabilities than far misses (*Figure 8*D).

**Video 5.** Movie for a hit stimulus (outward model with multiple units, where *M* = 32). Top left panel: the same as in the top row of *Figure 8*A; bottom left, top right, bottom left panels: the same as in *Video 1* but with more units. The movie has been slowed down by a factor of 10.

**Video 6.** Movie for a hit stimulus (inward model with multiple units, where *M* = 32). The same arrangement as *Video 5* but for an inward model.

**Video 7.** Movie for a miss stimulus (outward model with multiple units, where *M* = 32). The same arrangement as *Video 5*.

**Video 8.** Movie for a miss stimulus (inward model with multiple units, where *M* = 32). The same arrangement as *Video 6*.

**Video 9.** Movie for a retreat stimulus (outward model with multiple units, where *M* = 32). The same arrangement as *Video 5*.

**Video 10.** Movie for a retreat stimulus (inward model with multiple units, where *M* = 32). The same arrangement as *Video 6*.

**Video 11.** Movie for a rotation stimulus (outward model with multiple units, where *M* = 32). The same arrangement as *Video 5*.

**Video 12.** Movie for a rotation stimulus (inward model with multiple units, where *M* = 32). The same arrangement as *Video 6*.

### Large populations of units improve performance and favor outward filters

Since a larger number of units will cover an increasing spatial area of the visual field, the population of units can in principle provide more information about the incoming signals. In general, the models perform better as the number of units *M* increases (*Figure 9*A). When *M* is above 32, both the ROC-AUC and PR-AUC scores are almost 1 (Methods and Materials), which indicates that the model is very accurate on the binary classification task presented by the four types of synthetic stimuli.

**Figure 9.**
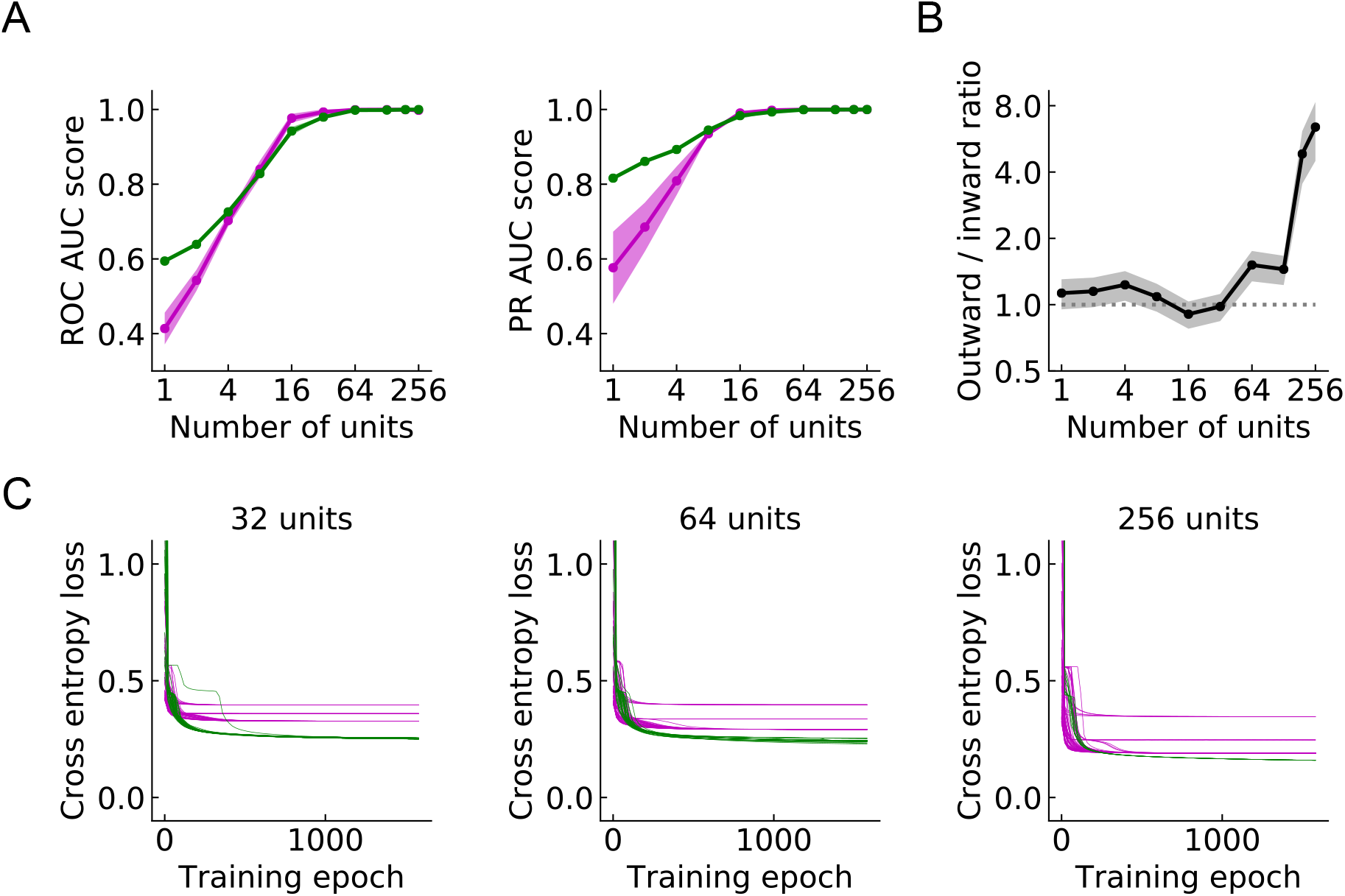
Large populations of units improve performances and favor outward solutions (Methods and Materials). (A) Both ROC and PR AUC scores increase as the number of units increases. Lines and dots: average scores; shading: one standard deviation of the scores over the trained models. Magenta: outward solutions; green: inward solutions. (B) The black line and dots show the ratio of the numbers of the two types of the solutions in the set of randomly initialized, trained models. The grey shading is one standard deviation, assuming that the distribution is binomial (Methods and Materials). The dotted horizontal line indicates the ratio of 1. (C) As the population of units increases, cross entropy losses of the outward solutions approach the losses of the inward solutions.

In addition, we calculated the ratio of the number of outward filters to inward filters that arise out of 200 random initializations in models with *M* units, as we swept *M*. Interestingly, as the number of units increases, an increasing proportion of solutions have outward filters (*Figure 9*B). For models with 256 units, the chance that an outward filter appears as a solution is almost 90% compared with the roughly 50% when *M* = 1. As *M* increases, the outward solutions become closer to the inward ones on both AUC scores and cross entropy losses (*Figure 9*A, C). These results suggest that as more units are added, optimization favors the outward solutions.

### Activation patterns of computational solutions resemble biological responses

The outward solutions have a receptive field structure that is similar to LPLC2 neurons, based on their anatomy and functional studies. However, it is not clear whether these models possess the functional properties of LPLC2 neurons, which have been studied systematically [Klapoetke et al., 2017, Von Reyn et al., 2017, Ache et al., 2019]. To see how trained units compare to LPLC2 neuron properties, we presented stimuli to the trained model (*Figure 10*A) to compare its responses to those measured in LPLC2 to similar stimuli.

**Figure 10.**
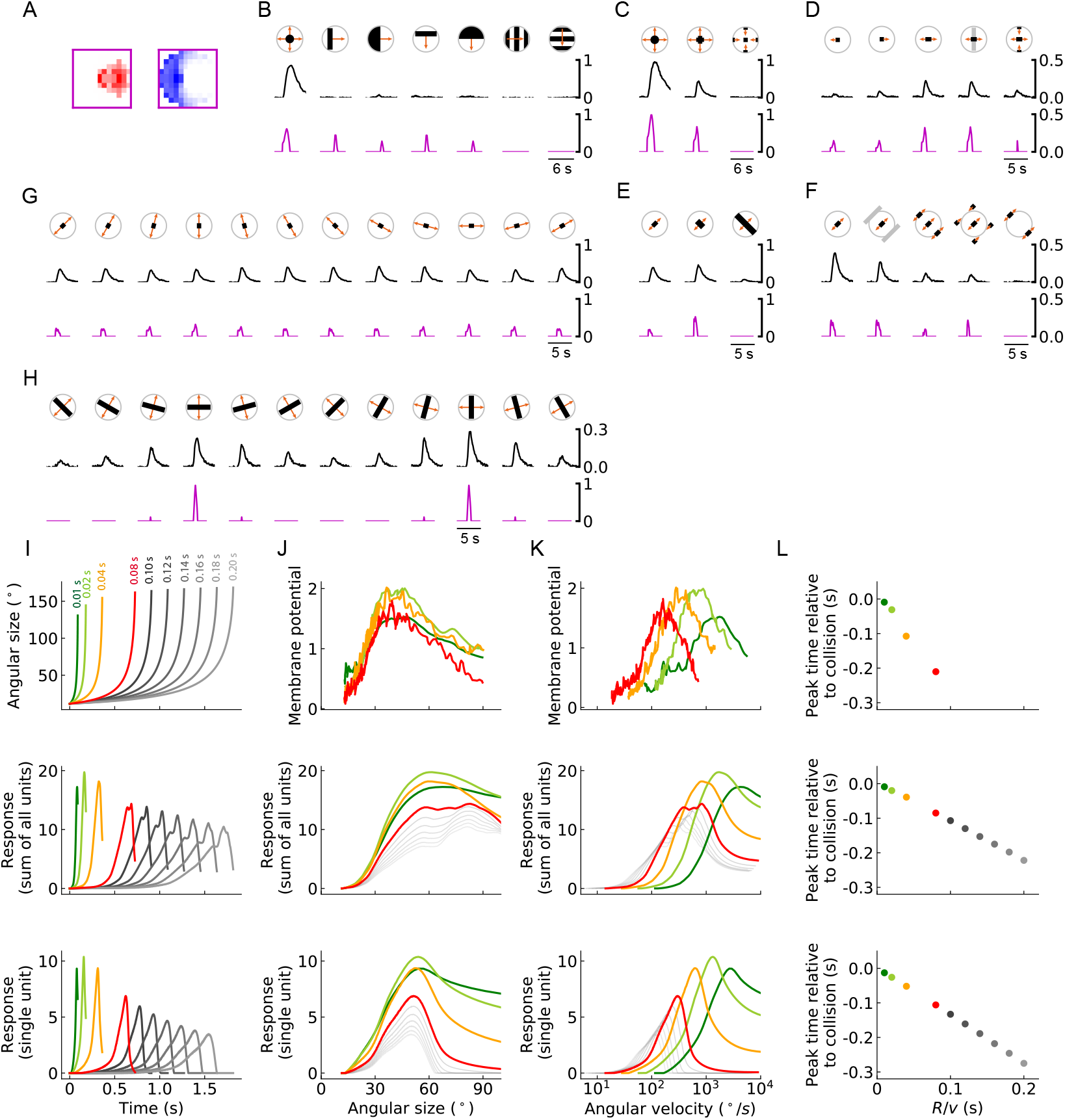
Models trained on binary classification tasks exhibit similar responses to LPLC2 neurons observed in experiments. (A) Excitatory and inhibitory filters of an outward solution with 256 units. (B-H) Comparisons of the responses of the solution in (A) and LPLC2 neurons to a variety of stimuli (Methods and Materials). Black lines: data [Klapoetke et al., 2017]; magenta lines: model. Compared with the original plots [Klapoetke et al., 2017], all the stimuli icons here except the ones in (B) have been rotated 45 degrees to match the cardinal directions of LP layers as described in this study. (I) Top: temporal trajectories of the angular sizes for different *R/v* ratios (color labels apply throughout (I-L)) (Methods and Materials). Middle: response as a function of time for the sum of all 256 units. Bottom: response as a function of time for one of the 256 units. (J-L) Top: experime newtal data (LPLC2/non-LC4 components of GF activity. Data from [Von Reyn et al., 2017, Ache et al., 2019]). Middle: sum of all 256 units. Bottom: response of one of the 256 units. Responses as function of angular size (J), response as function of angular velocity (K), relationship between peak time relative to collision and *R/v* ratios (L). We considered the first peak when there were two peaks in the response, such as in the grey curves in the middle panel of (I). **Figure 10–Figure supplement 1.** Same as in the main figure but for a different outward model obtained from the same training procedure.

The model behaves similarly to LPLC2 neurons on many different types of stimuli. Not surprisingly, the model is selective to loom signals and does not have strong responses to non-looming signals (*Figure 10*B). Moreover, the model closely follows the response of LPLC2 neurons to various expanding bar stimuli, including the inhibitory effects of inward motion (*Figure 10*C, D). In addition, in experiments, motion signals that appear at the periphery of the receptive field suppress the activity of the LPLC2 neurons (periphery inhibition) [Klapoetke et al., 2017], and this phenomenon is successfully predicted by the model (*Figure 10*E, F) due to the broad inhibitory filters that the model learns (*Figure 10*A). Interestingly, the model also correctly predicts response patterns of the LPLC2 neurons for expanding bars with different orientations (*Figure 10*G, H).

The ratio of object size to approach velocity, or *R/v*, is an important parameter for looming stimuli, and many studies have investigated how the response patterns of loom-sensitive neurons depend on this ratio (Top panels in *Figure 10*I, J, K, L) [Gabbiani et al., 1999, Von Reyn et al., 2017, Ache et al., 2019, De Vries and Clandinin, 2012]. Here, we presented the trained model (*Figure 10*A) with hit stimuli with different *R/v* ratios, and compared its behaviors with the experimental data (*Figure 10*I-L)). Surprisingly, although our model only has the angular velocities as the inputs (*Figure 3*), it reliably encodes the angular sizes rather than the angular velocities, indicated by the collapsed response curves (up to different scales) when plotted against the angular sizes (*Figure 10*J) [Von Reyn et al., 2017], though the model curve shapes do not exactly match the experimental ones. On the contrary, for angular velocities, the response curves shift for different *R/v* ratios, which means they depend on the velocities *v* of the object (*R* is fixed to be 1). Both of these response properties are consistent with properties of LPLC2. Meanwhile, a canonical linear relationship between the peak response time relative to the collision and the *R/v* ratio is also reproduced by the optimized model (*Figure 10*L) [Gabbiani et al., 1999, Ache et al., 2019].

Importantly, a different outward solution from the same training procedure could reproduce many of the same effects (*Figure 10–Figure Supplement 1*), but it predicts the patterns in the wide expanding bars differently and out of phase from the biological data (*Figure 10–Figure Supplement 1*H). This different solution also does a poor job predicting the response curves of the LPLC2 neurons to looming signals with different *R/v* ratios, although the collapsed and shifted features remain when plotted as functions of angular size and velocity (*Figure 10–Figure Supplement 1*J, K). This shows that even within the family of learned outward solutions, there is variability in the learned response properties. Though solving the inference problem obtains many of the response properties, additional constraints would be required to more precisely reproduce the LPLC2 responses.

## Discussion

In this study, we have shown that training a simple network to detect collisions gives rise to a computation that closely resembles neurons that are sensitive to looming signals. Specifically, we optimized a neural network model to detect whether an object is on a collision course based on the visual motion signals (*Figure 3*), and found that one class of optimized solution matched the anatomy of motion inputs to LPLC2 neurons (*Figure 1, Figure 5, Figure 6*). Importantly, this solution can reproduce a large range of experimental observations of LPLC2 neuron responses (*Figure 10*) [Klapoetke et al., 2017, Von Reyn et al., 2017, Ache et al., 2019].]

The radially structured dendrites of the LPLC2 neuron in the lobula plate can account for its response to motion radiating outward from the receptive field center [Klapoetke et al., 2017]. Our results show that the logic of this computation can be understood in terms of inferential loom detection by the *population* of units. In particular, for an individual detector unit, an inward structure makes a better loom detector than an outward structure, since it is sensitive to colliding objects originating from a wider array of incoming angles (*Figure 7*). As the number of units across visual space increases, the outward-sensitive receptive field structure is represented more often in the optimal solution. The solution depends on the number of detectors, and this is likely related to the increasing overlap in receptive fields as the population grows (*Figure 6*). This result is consistent with prior work showing that populations of neurons often exhibit different and improved coding strategies compared to individual neurons [Pasupathy and Connor, 2002, Georgopoulos et al., 1986, Vogels, 1990, Franke et al., 2016, Zylberberg et al., 2016, Cafaro et al., 2020]. Thus, understanding anatomical, physiological, and algorithmic properties of individual neurons can require considering the population response. The solutions we found to the loom inference problem suggest that individual LPLC2 responses should be interpreted in light of the population of LPLC2 responses.

Surprisingly, the trained outward solutions exhibits the properties of an angular size encoder (*Figure 10*I-L), even though the inputs to the model are a field of motion signals. There are two ways that this tuning arises. First, in a hit stimulus, the angular size and angular velocity are strongly correlated [Gabbiani et al., 1999], which means the angular size affects the magnitude of the motion signals. Second, the angular size is proportional to the length of the outward-moving edges of hitting objects. The angular circumference of the hit stimulus determines how many motion detectors are activated, so that integrated motion signal strength is related to the size. Both of these effects influence the response patterns of the model units (and the LPLC2 neurons).

Our results shed light on discussions of *η*-like (encoding angular size) and *ρ*-like (encoding angular velocity) looming sensitive neurons in the literature [Gabbiani et al., 1999, Wu et al., 2005, Liu et al., 2011, Shang et al., 2015, Temizer et al., 2015, Dunn et al., 2016, Von Reyn et al., 2017, Ache et al., 2019]. In particular, these optimized models clarify an interesting but puzzling fact: LPLC2 neurons transform their inputs of direction-selective motion signals to computations of angular size [Ache et al., 2019]. Consistently, our model shows the linear relationship between the peak time relative to collision and the *R/v* ratio, which looming sensitive neurons that encode angular size should follow [Peek and Card, 2016]. In both cases, these properties appear to be the simple result of training the constrained model to reliably detect looming stimuli.

The units of the outward solution exhibit sparsity in their responses to looming stimuli, in contrast to the denser representations in the inward solution (*Figure 8*). During a looming event, most of the units are quiet and only a few adjacent units have very large activities, reminiscent of sparse codes that seem to be favored, for instance, in cortical encoding of visual scenes [Olshausen and Field, 1996, 1997]. Since the readout of our model is a summation of the activities of the units, sparsity does not directly affect the performance of the model, but is an attribute of the favored solution. For a model with a different loss function or noise, the degree of sparsity might be crucial. For instance, the sparse code of the outward model might make it easier to localize the hit stimulus [Morimoto et al., 2020], or might make the population response more robust to noise [Field, 1994].

Experiments have shown that inhibitory circuits play an important role for the selectivity of LPLC2 neurons. For example, motion signals at the periphery of the receptive field of an LPLC2 neuron inhibit its activity; such peripheral inhibition causes various interesting response patterns of the LPLC2 neurons to different types of stimuli [Klapoetke et al., 2017]. However, the structure of this inhibitory field is not fully understood, and our model provides a tool to investigate how the inhibitory inputs to LPLC2 neurons affect circuit performance on loom detection tasks. Specifically, strong inhibition on the periphery of the receptive field arises naturally in the outward solutions after optimization [Klapoetke et al., 2017]. The broad inhibition appears in our model to suppress responses to the non-hit stimuli. As in the data, the inhibition is broader than one might expect if the neuron were simply being inhibited by inward motion.

The synthetic stimuli used to train models in this study were unnatural in two ways. The first way was in the proportion of hits and non-hits. We trained with 25% of the training data representing hits. The true fraction of hits among all stimuli encountered by a fly is undoubtedly much less, and this affects how the loss function weights different types of errors. It is also clear that a false-positive hit (in which a fly might jump to escape an object not on collision course) is much less penalized during evolution than a false-negative (in which a fly doesn’t jump and an object collides, presumably to the detriment of the fly). It remains unclear how to choose these weights in the training data or in the loss function, but they affect the receptive field weights optimized by the model.

The second issue with the stimuli is that they were caricatures of stimulus types, but did not incorporate the richness of natural stimuli. This richness could include natural textures and spatial statistics [Ruderman and Bialek, 1994], which seem to impact motion detection algorithms [Fitzgerald and Clark, 2015, Leonhardt et al., 2016, Chen et al., 2019]. This richness could also include more natural trajectories for approaching objects. Another way to enrich the stimuli would be to add noise, either in inputs to the model or in the model’s units themselves. These aspects of the stimuli were all neglected in this initial study, in part because it is difficult to find characterizations of natural looming events. An interesting future direction will be to investigate the effects of more complex and naturalistic stimuli on the model’s filters and performance, as well as on LPLC2 neuron responses themselves.

For simplicity, this model did not impose the hexagonal geometry of the compound eye ommatidia. Instead, we assume that the visual field is separated into a Cartesian lattice with 5° spacing, each representing a local motion detector with two spatially separated inputs (*Figure 3*). This simplification alters slightly the geometry of the motion signals compared to the real motion detector receptive fields [Shinomiya et al., 2019]. This could potentially affect the learned spatial weightings and reproduction of the LPLC2 responses to various stimuli, since the specific shapes of the filters matter (*Figure 10*). Thus, the hexagonal ommatidial structure and the full extent of inputs to T4 and T5 might be crucial if one wants to make comparisons with the dynamics and detailed responses of LPLC2 neurons. However, this geometric distinction seems unlikely to affect the main results of how to infer the presence of hit stimuli.

Our model requires a field of estimates of the local motion. Here, we used the simplest model – the Hassenstein-Reichardt correlator model *Equation 3* (Methods and Materials) [Hassenstein and Reichardt, 1956] – but the model could be extended by replacing it with a more sophisticated model for motion estimation. Some biophysically realistic ones might take into account synaptic conductances [Gruntman et al., 2018, 2019, Badwan et al., 2019, Zavatone-Veth et al., 2020]. Alternatively, in natural environments, contrasts fluctuate in time and space. Thus, if one includes more naturalistic spatial and temporal patterns, one might consider a motion detection model that can adapt to changing contrasts in time and space [Drews et al., 2020, Matulis et al., 2020].

Our neural network model is highly constrained by the specific anatomy of LPLC2 circuits, and no unnecessary layers were added. The resulting model is a shallow neural network (*Figure 1* and *Figure 4*. This shallowness leads to limited dimensionality of the model, which will be prone to finding non-optimal local minima during training. Indeed, in many cases, the training resulted in models with poor performance and filters with weights very close to zero. We minimized this problem by choosing the initialization scales for the filters so that optimization resulted in meaningful models with structured filters about half of the time. However, the ratio of the outward and inward solutions (*Figure 9*B) was not affected by the initialization scales.

Although the outward filter of the unit emerges naturally from our gradient descent training protocol, that does not mean that the structure is learned by LPLC2 neurons in the fly. There is some experience dependent plasticity in the fly eye [Kikuchi et al., 2012], but these visual computations are likely to be primarily genetically determined. Thus, one could think of the computation of the LPLC2 neuron as being shaped through millions of years of evolution. Interestingly, optimization algorithms similar to evolution may be able to avoid getting stuck in local optima [Stanley et al., 2019], and thus work well with the sort of shallow neural network found in the fly eye.

In this study, we focused on the motion signal inputs to LPLC2 neurons, and we neglected other inputs to LPLC2 neurons, such as inputs coming from the lobula that likely report non-motion visual features. It would be interesting to investigate how this additional non-motion information would affect the performance and optimal solutions of the inference units. For instance, another lobula columnar neurons, LC4, is loom sensitive and receives inputs in the lobula [Von Reyn et al., 2017]. The LPLC2 and LC4 neurons are the primary excitatory inputs to the GF, which mediates the escape behavior of a fly [Von Reyn et al., 2014, Ache et al., 2019]. The inference framework set out here would allow one to incorporate of parallel non-motion intensity channels, either by adding them into the inputs to the LPLC2-like units, or by adding in a parallel population of LC4-like units. This would require a reformulation of the probabilistic model in *Equation 5*. Notably, one of the most studied loom detecting neurons, the lobula giant movement detector (LGMD) in locusts, does not appear to receive direction-selective inputs, as LPLC2 does [Rind and Bramwell, 1996, Gabbiani et al., 1999]. Thus, the inference framework set out here can be flexibly modified to investigate loom detection under a wide variety of constraints and inputs, which allow it to be applied to other neurons, beyond LPLC2.

## Methods and Materials

### Code availability

Code to perform all simulations in this paper and to reproduce all figures is available at http://www.github.com/ClarkLabCode/LoomDetectionANN.

### Coordinate system and stimuli

We designed a suite of visual stimuli to simulate looming objects, retreating objects, and rotational visual fields. In this section, we describe the suite of stimuli and the coordinate systems used in our simulations (*Figure 4–Figure Supplement 1*).

In our simulations and training, the fly is at rest on a horizontal plane, with its head pointing in a specific direction. The fly head is modeled to be a point particle with no volume. A three dimensional right-handed frame of reference Σ is set up and attached to the fly head at the origin. The *z* axis points in the anterior direction from the fly head, perpendicular to the line that connects the two eyes, and in the horizontal plane of the fly; the *y* axis points toward the right eye, also in the horizontal plane; and the *x* axis points upward and perpendicular to the horizontal plane. Looming or retreating objects are represented in this space by a sphere with radius *R* =1, and the coordinates of an object’s center at time *t* are denoted as **r**(*t*) = (*x*(*t*), *y*(*t*), *z*(*t*)). Thus, the distance between the object center and the fly head is 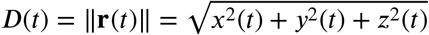.

Within this coordinate system, we set up cones to represent individual units. The receptive field of LPLC2 neurons is measured at roughly 60° in diameter [Klapoetke et al., 2017]. Thus, we here model each unit as a cone with its vertex at the origin and with half-angle of 30°. For each unit *m* (*m* = 1, 2,…, *M*), we set up a local frame of reference Σ_*m*_ (*Figure 4–Figure Supplement 1*): the *z_m_* axis is the axis of the cone and its positive direction points outward from the origin. The local Σ_*m*_ can be obtained from Σ by two rotations: around *x* of Σ and around the new *y*′ after the rotation around *x*. For each unit, its cardinal directions are defined as: upward (positive direction of *x_m_*), downward (negative direction of *x_m_*), leftward (negative direction of *y_m_*) and rightward (positive direction of *y_m_*). To get the signals that are received by a specific unit *m*, the coordinates of the object in Σ are rotated to the local frame of reference Σ_*m*_.

Within this coordinate system, we can set up cones representing the extent of a spherical object moving in the space. The visible outline of a spherical object spans a cone with its point at the origin. The half-angle of this cone is a function of time and can be denoted as *θ*_s_(*t*):

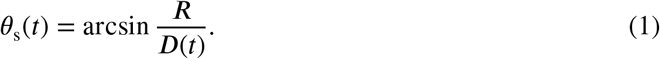

One can calculate how the cone of the object overlaps with the receptive field cones of each unit.

There are multiple layers in the fly visual system [Takemura et al., 2017], but here we focus on two coarse grained stages of processing: (1) the estimation of local motion direction from optical intensities by motion detection neurons T4 and T5 and (2) the integration of the flow fields by LPLC2 neurons. In our simulations, the interior of the *m*th unit cone is represented by a *N*-by-*N* matrix, so that each element in this matrix (except the ones at the four corners) indicates a specific direction in the angular space within the unit cone. If an element also falls within the object cone, then its value is set to 1; otherwise it is 0. Thus, at each time *t*, this matrix is an optical contrast signal and can be represented by *C*(*x_m_, y_m_, t*), where (*x_m_, y_m_*) are the coordinates in Σ_*m*_. In general, *N* should be large enough to provide good angular resolutions. Then, *K*^2^ (*K* < *N*) motion detectors are evenly distributed within the unit cone, with each occupying an *L*-by-*L* grid in the *N*-by-*N* matrix, where *L* = *N/K*. This *L*-by-*L* grid represents a 5°-by-5° square in the angular space, consistent with the approximate spacing of the inputs of motion detectors T4 and T5. This arrangement effectively upsamples the spatial resolution of the intensity data before it is discretized into motion signals with a resolution of 5°. Since the receptive field of an LPLC2 neuron is roughly 60°, the value of *K* is chosen to be 12. To get sufficient angular resolution for the local motion detectors, *L* is set to be 4, so that *N* is set to 48.

Each motion detector is assumed to be a Hassenstein Reichardt Correlator (HRC) and calculates local flow fields from *C*(*x_m_, y_m_, t*) [Hassenstein and Reichardt, 1956, Potters and Bialek, 1994]. The HRC used here has two inputs, separated by 5° in angular space. Each input applies first a spatial filter on the contrast *C*(*x_m_, y_m_, t*) and then temporal filters:

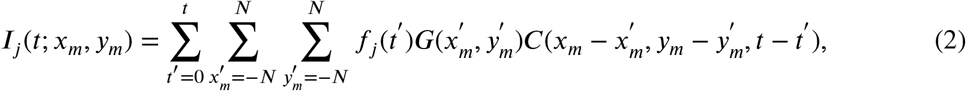

where *f_j_*(*j* ∈ 1, 2) is a temporal filter and *G* is a discrete 2d Gaussian kernel with mean 0° and standard deviation of 2.5° to approximate the acceptance angle of the fly photoreceptors [Stavenga, 2003]. The temporal filter *f*_1_ was chosen to be an exponential function *f*_1_(*t*′) = (1/*τ*) exp(−*t*′ /*τ*) with *τ* set to 0.03 seconds [Salazar-Gatzimas et al., 2016], and *f*_2_ a delta function *f*_2_ = *δ*(*t*′). This leads to

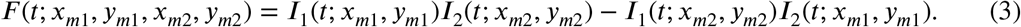

as the local flow field at time *t* between two inputs located at (*x*_*m*1_, *y*_*m*1_) and (*x*_*m*2_, *y*_*m*2_).

Four types of T4 and T5 neurons have been found that project to layers 1, 2, 3, and 4 of the lobula plate. Each type is sensitive to one of the cardinal directions: down, up, left, right [Maisak et al., 2013]. Thus, in our model, there are four non-negative, local flow fields that serve as the only inputs to the model: *U*_−_(*t*) (downward, corresponding LP layer 4), *U*_+_(*t*) (upward, LP layer 3), *V*_−_(*t*) (leftward, LP layer 1) and *V*_+_(*t*) (rightward, LP layer 2), each of which is a *K*-by-*K* matrix. To calculate these matrices, two sets of motion detectors are needed, one for the vertical directions and one for the horizontal directions. The HRC model in *Equation 3* is direction sensitive and is opponent, meaning that for motion in the preferred (null) direction, the output of the HRC model is positive (negative). Thus, assuming that upward (rightward) is the preferred vertical (horizontal) direction, we obtain the non-negative elements of the four flow fields as

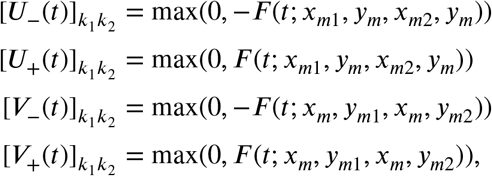

where *k*_1_, *k*_2_ ∈ {1, 2,…, *K*} and |·| represents the absolute value. In the above expressions, it implies, for [*U*_−_(*t*)]_*k*_1_*k*_2__ and [*U*_+_(*t*)]_*k*_1_*k*_2__, the vertical motion detector at (*k*_1_, *k*_2_) has its two inputs located at (*x*_*m*1_, *y_m_*) and (*x*_*m*2_, *y_m_*), respectively. Similarly, for for [*V*_−_(*t*)]_*k*_1_*k*_2__ and [*V*_+_(*t*)]_*k*_1_*k*_2__, the horizontal motion detector at (*k*_1_, *k*_2_) has its two inputs located at (*x_m_*, *y*_*m*1_) and (*x_m_*, *y*_*m*2_). Using the opponent HRC output as the motion signals for each layer is reasonable because the motion detectors T4 and T5 are highly direction-selective over a large range of inputs [Maisak et al., 2013, Creamer et al., 2018] and synaptic, 3-input models for T4 are approximately equivalent to opponent HRC models [Zavatone-Veth et al., 2020].

We simulated the trajectories **r**(*t*) of the object in the frame of reference Σ at a time resolution of 0.01 seconds. For hit, miss, and retreat cases, the trajectories of the object are always straight lines (i.e., ballistic motion), and the velocities of the object were randomly sampled from a range [2*R*, 10*R*](*s*^−1^) with the trajectories confined to be within a sphere of 5*R* centered at the fly head. The radius of the object, *R*, is always set to be 1 except in the rotational stimuli. To generate rotational stimuli, we placed 100 objects with various radii randomly selected from [0, 1] at random distances ([5, 15]) and positions around the fly, and rotated them all around a randomly chosen axis. The rotational speed was chosen from a Gaussian distribution with mean 0°/*s* and standard deviation 200°/*s*, a reasonable rotational velocity for walking flies [DeAngelis et al., 2019].

We reproduced a range of stimuli used in a previous study [Klapoetke et al., 2017] and tested our trained model on them (*Figure 10*B-H). To match the cardinal directions of LP layers (*Figure 1*), we have rotated the stimuli (except in *Figure 10*B) 45 degrees compared with the ones displayed in the figures in [Klapoetke et al., 2017]. The disc (*Figure 10*B, C) expands from 20° to 60° with an edge speed of 10°/*s*. All the bar and edge motions have an edge speed of 20°/*s*. The width of the bars are 60° (right panel of *Figure 10*E, and *Figure 10*H), 20° (middle panel of *Figure 10*E), and 10° (all the rest). All the responses of the models (except in *Figure 10*B) have been normalized by the peak of the response to the expanding disc (*Figure 10*B).

We created a range of hit stimuli with various *R/v* ratios: 0.01 *s*, 0.02 *s*, 0.04 *s*, 0.08 *s*, 0.10 *s*, 0.12 *s*, 0.14 *s*, 0.16 *s*, 0.18 *s*, 0.20 *s*. The radius *R* of the spherical object is fixed to be 1, and the velocity is changed accordingly to achieve different *R/v* ratios.

### Models

Experiments have shown that an LPLC2 neuron has four dendritic structures in the four LP layers, and that they receive direct excitatory inputs from T4/T5 motion detection neurons [Maisak et al., 2013, Klapoetke et al., 2017]. It has been proposed that each dendritic structure also receives inhibitory inputs mediated by lobulate plate intrinsic interneurons, such as LPi4-3 [Klapoetke et al., 2017]. Accordingly, our models have two types of nonnegative filters, one excitatory and one inhibitory (*Figure 4*, represented by *W*^e^ and *W*^i^, respectively. Each filter is a 12-by-12 matrix. We rotate *W*^e^ and *W*^i^ counterclockwise by multiples of 90° to obtain the filters that are used to integrate the four motion signals: *U*_−_(*t*), *U*_+_(*t*), *V*_−_(*t*), *V*_+_(*t*). Specifically, we define the corresponding four excitatory filters as: 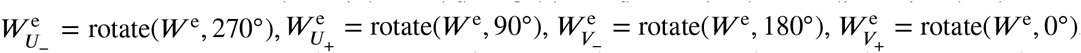, and the inhibitory filters as: 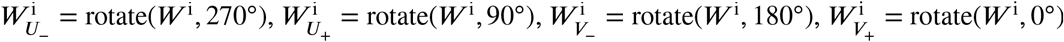. In addition, we impose mirror symmetry to the filters, and with the above definitions of the rotated filters, the upper half of *W*^e^ is a mirror image of the lower half of *W*^e^. The same mirror symmetry applies to *W*^i^. Thus, there are in total 144 parameters in the two sets of filters. In fact, since only the elements within a 60 degree cone contribute to the filter for the units, the corners are excluded, resulting in only 112 trainable parameters in the excitatory and inhibitory filters.

In computer simulations, the weights and flow fields are flattened to be one-dimensional column vectors. The responses of the inhibitory units are:

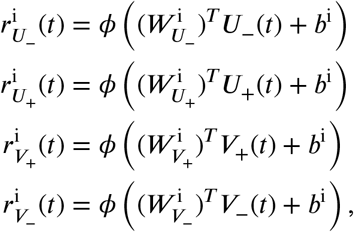

where *ϕ*(·) = max(·, 0) is the rectified linear activation function, and 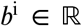 is the intercept. The response of a single unit *m* is

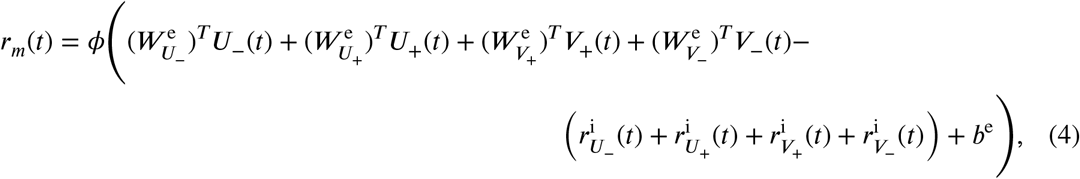

where 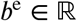 is the intercept (*Figure 4*). The inferred probability of hit for a specific trajectory is

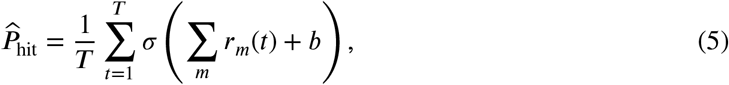

where *T* is the total number of time steps in the trajectory and *σ*(·) is the sigmoid function. Since we are adding three intercepts *b*^i^, *b*^e^, and *b*, there are 115 parameters to train in this model.

### Training and testing

We created a synthetic data set containing four types of motion: *loom-and-hit, loom-and-miss, retreat*, and *rotation*. The proportions of these types were 0.25, 0.125, 0.125, and 0.5, respectively. In total, there were 5200 trajectories, with 4,000 for training and 1,200 for testing. Trajectories with motion type *loom-and-hit* are labeled as hit or *y_n_* = 1 (probability of hit is 1), while trajectories of other motion types are labeled as non-hit or *y_n_* = 0 (probability of hit is 0), where *n* is the index of each specific sample. Models with smaller *M* have fewer trajectories in the receptive field of any unit. For stability of training, we therefore increased the number of trajectories by factors of eight, four, and two for *M* = 1,2,4, respectively.

The loss function to be minimized in our training was the cross entropy between the label *y_n_* and the inferred probability of hit 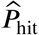, and averaged across all samples, together with a regularization term:

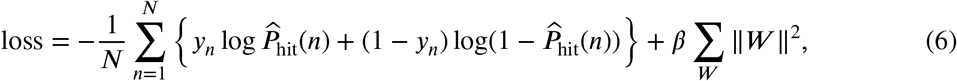

where 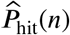 is the inferred probability of hit for sample *n*, *β* is the strength of the *ℓ*_2_ regularization, and *W* represents all the effective parameters in the two excitatory and inhibitory filters.

The strength of the regularization *β* was set to be 10^−4^, which was obtained by gradually increasing *β* until the performance of the model on test data started to drop. The regularization sped up convergence of solutions, but the regularization strength did not strongly influence the main results in the paper.

To speed up training, rather than taking a temporal average as shown in *Equation 5*, a snapshot was sampled randomly from each trajectory, and the probability of hit of this snapshot was used to represent the whole trajectory, i.e., 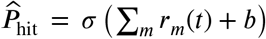, where *t* is a random sample from {1,2,…, *T*}. Mini-batch gradient descent was used in training, and the learning rate was 0.001.

After training, the models were tested on the entire trajectories with the probability of hit defined in *Equation 5*. Models trained only on snapshots performed well on the test data. During testing, the performance of the model was evaluated by the area under the curve (AUC) of the receiver operating characteristic (ROC) and precision-recall (PR) curves [Hanley and McNeil, 1982, Davis and Goadrich, 2006]. TensorFlow [Abadi et al., 2016] was used to train all models.

### Clustering the solutions

We used the following procedure to cluster the solutions. Each solution had an excitatory and an inhibitory filter. We flattened these two filters, and concatenated them into a single vector. (The elements at the corners were deleted since they are outside of the receptive field.) Thus, each solution was represented by a vector, from which we calculated the cosine distance for each pair of solutions. The obtained distance matrix was then fed into a hierarchical clustering algorithm [Virtanen et al., 2020]. After obtaining the hierarchical clustering, the outward and inward filters were identified by their shape. We counted the non-zero filter elements corresponding to flow fields with components radiating outward and subtracted the number of non-zero filter elements corresponding to flow fields with components directed inward. If the resulted value was positive, the filters were labeled as outward; otherwise, the filters were labeled as inward. If the elements in the concatenated vector were all close to zero, then the corresponding filters were labeled as unstructured.

### Statistics

To calculate the fraction of active units for the model with *M* = 256 (*Figure 8*B), we looked at the response curves of each unit to all trajectories of a specific type of stimuli, and if the unit response is above the baseline (dotted lines in *Figure 7*B), then the unit is counted as active. So, for each trajectory/stimulus, we obtained the number of active units. After this, we calculated the mean and standard deviation across all the trajectories within each type of stimuli (hit, miss, retreat, rotation).

For a model with *M* units, where *M* ∈ {1,2,4, 8,16,32,64,128,192,256}, 200 random initializations were used to train it. Within these 200 training runs, the number of outward solutions *N*_out_ were (starting from smaller values of *M*) 44, 46, 48, 50, 48, 50, 53, 55, 58, 64, and the number of inward solutions *N*_in_ were 39, 40, 39, 46, 53, 51, 35, 38, 12, 10. The average score curves and dots in *Figure 9A* were obtained by taking the average among each type of solution, with the shading indicating two standard deviations. The curve and dots in *Figure 9*B are the ratio of the number of outward solutions to the number of inward solutions. To obtain error bars (grey shade), we considered the training results as a binomial distribution, with the probability of obtaining an outward solution being *N*_out_/(*N*_out_ + *N*_in_), and with the probability of obtaining an inward solution being *N*_in_/(*N*_out_ + *N*_in_). Thus, the standard deviation of this binomial distribution is 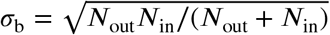. From this, we calculate the error bar as the propagated error [Morgan et al., 1990]:

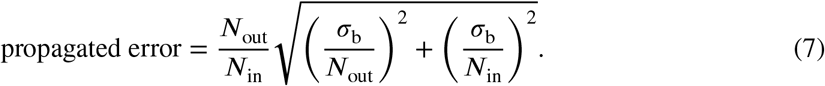

## Acknowledgments

Research supported in part by NSF grants DMS-1513594, CCF-1839308, DMS-2015397, a J.P. Morgan Faculty Research Award, and the Kavli Foundation. We thank G. Card and N. Klapoetke for sharing data traces from their paper. We thank members in the Clark laboratory for valuable discussions and comments.

---

**Figure 3–Figure supplement 1.**
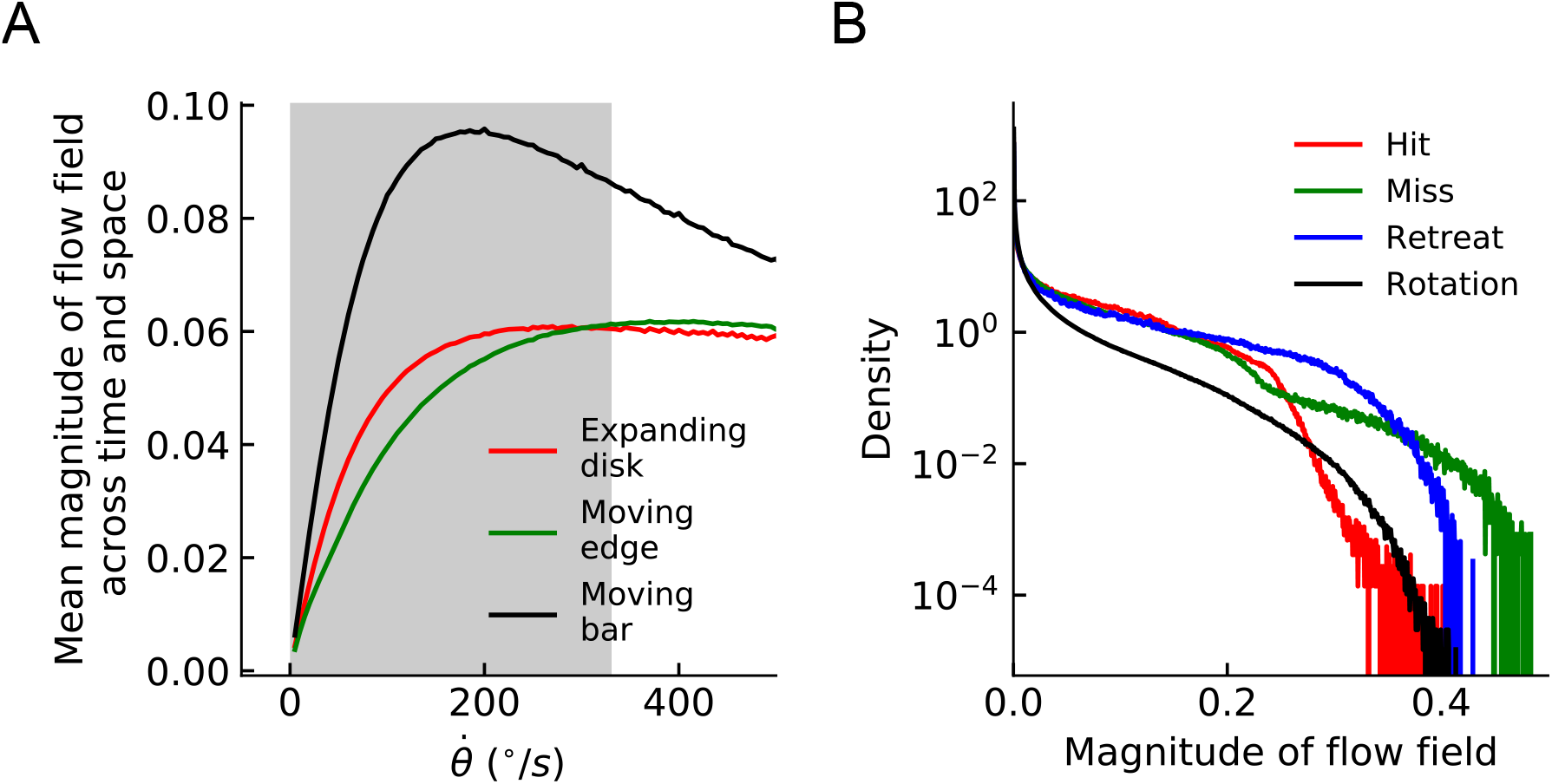
(A) Tuning curves of the HRC motion estimator for different stimuli. Grey shade indicates the velocity range used in simulations (Methods and Materials). (B) The distributions of the magnitude of all the estimated flow fields in the four cardinal directions for different types of stimuli.

**Figure 4–Figure supplement 1.**
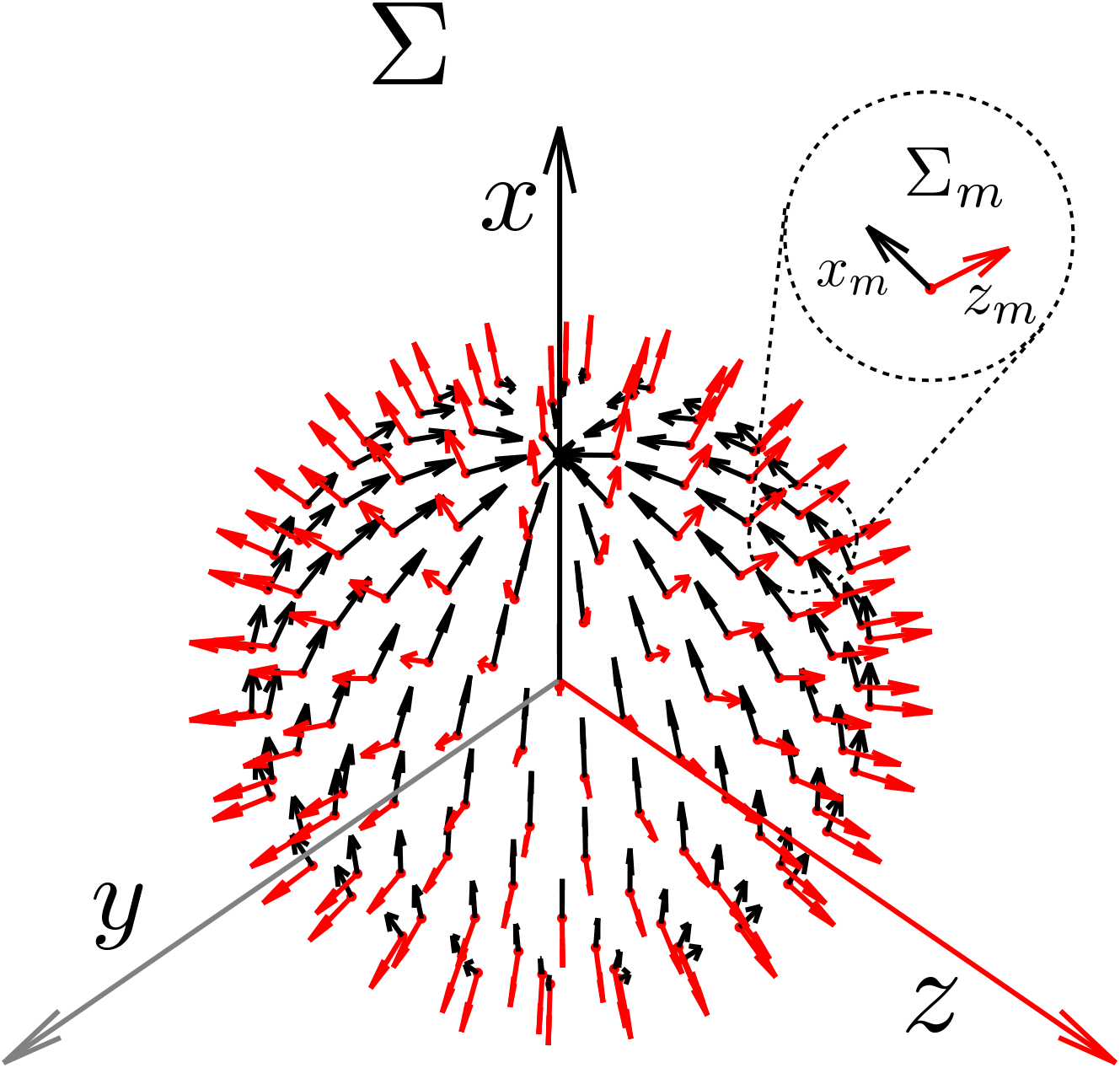
The coordinate system used in stimulus generation and modeling (Methods and Materials). The frame of reference Σ is fixed on the fly head. The frame of references Σ_*m*_ (*m* = 1,2,…, *M*) are associated with each local loom detector/model unit, the center of which is represented by a red dot. The origin of the Σ_*m*_ coincides with the origin of Σ, but for better illustration, we have translated its origin radially along *z_m_* so that all the Σ_*m*_’s sit on a sphere. For Σ_*m*_, only *x_m_* and *z_m_* are shown, and *y_m_* axis should be chosen such that Σ_*m*_ is right-handed. For the unit coordinate systems, *z_m_* is chosen to be normal to the unit sphere while *x_m_* points north tangent to the sphere. This system does not impose the left-right mirror symmetry of the fly eyes.

**Figure 5–Figure supplement 1.**
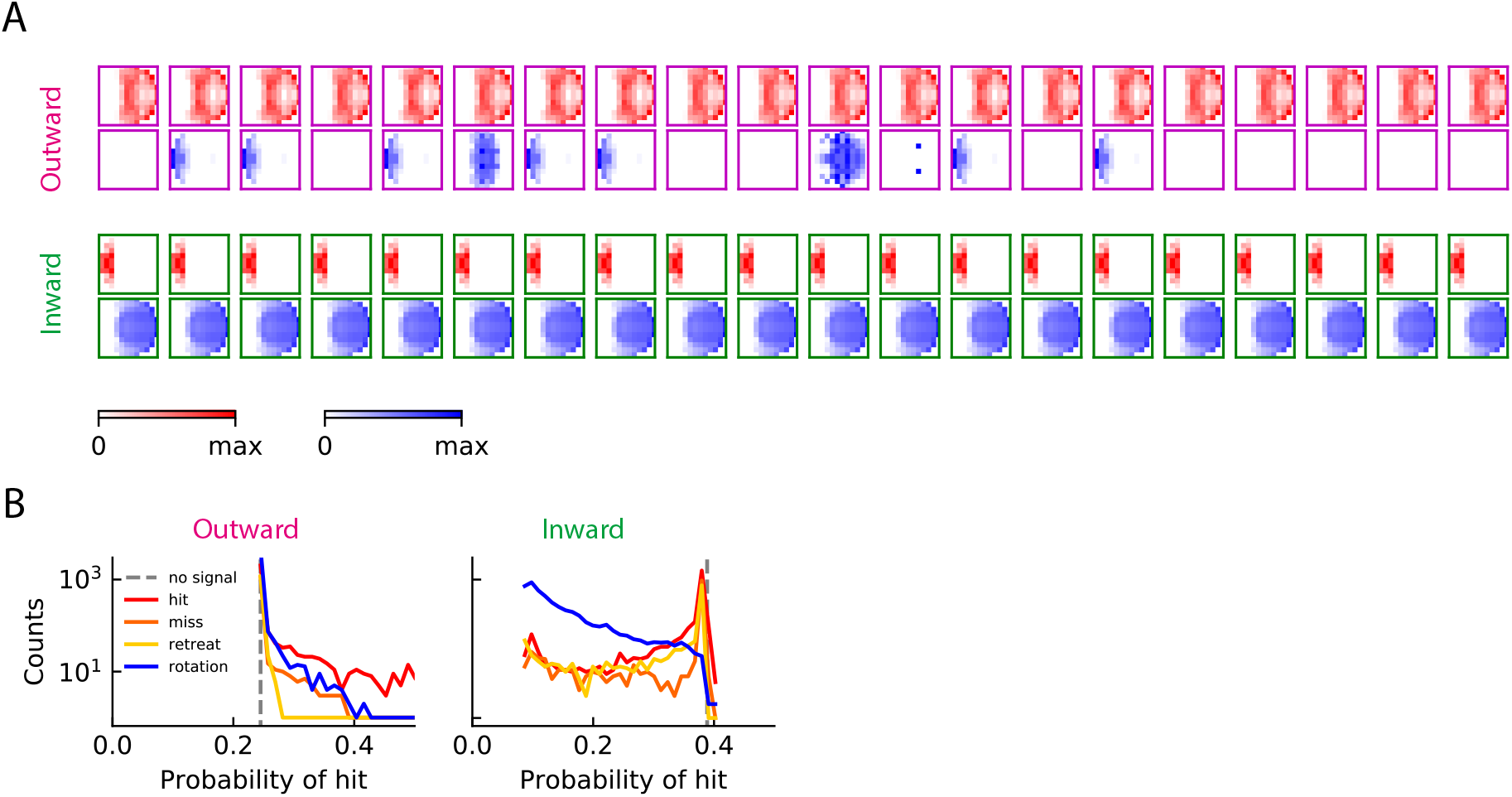
(A) Trained filters: outward solution (magenta) and inward solution (green). (B) Probability of hit inferred by a single unit for the four types of training stimuli.

**Figure 6–Figure supplement 1.**
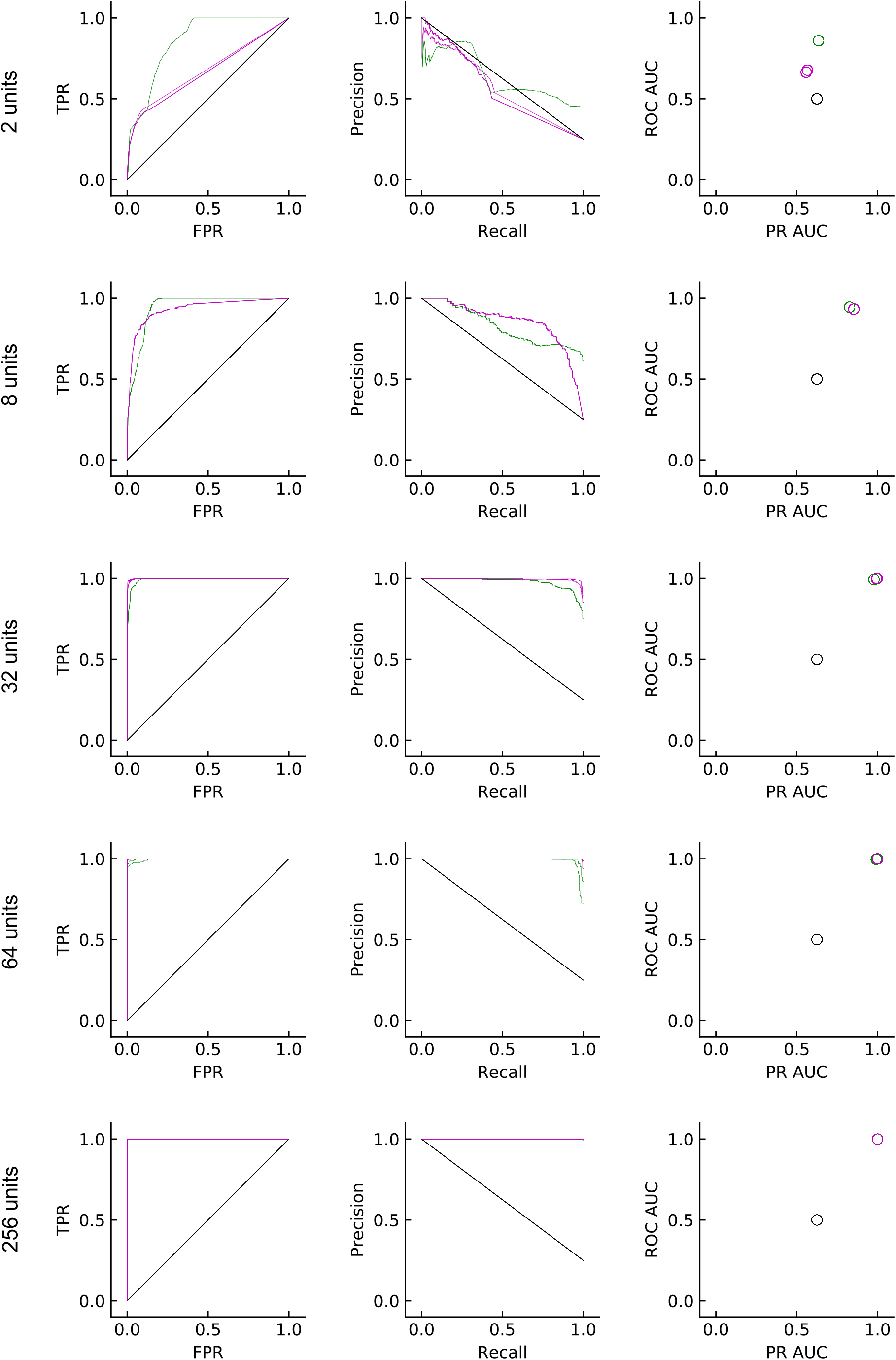
Same as in Figure 5D but for models with multiple units.

**Figure 6–Figure supplement 2.**
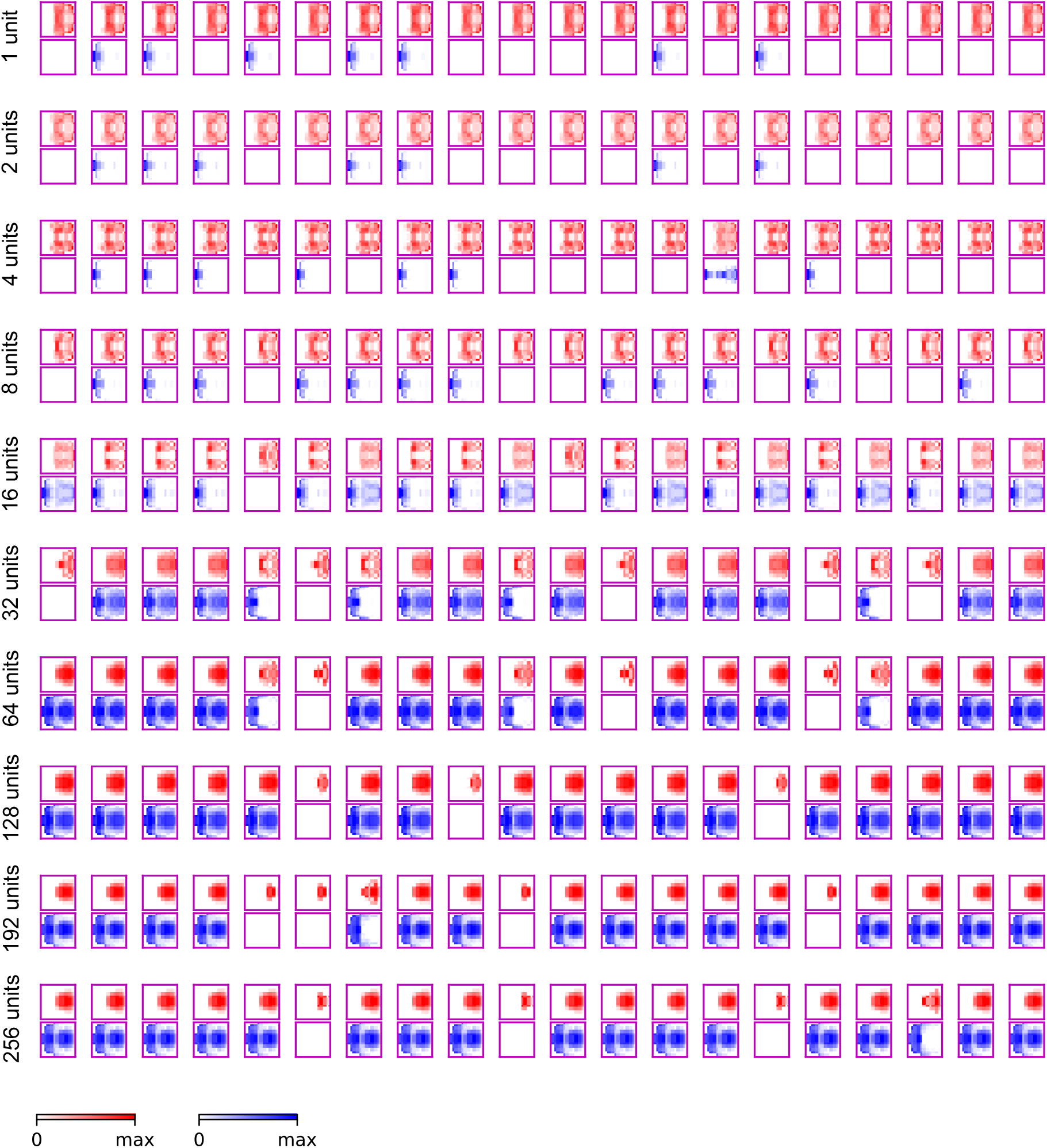
Outward solutions for models with different numbers of units.

**Figure 6–Figure supplement 3.**
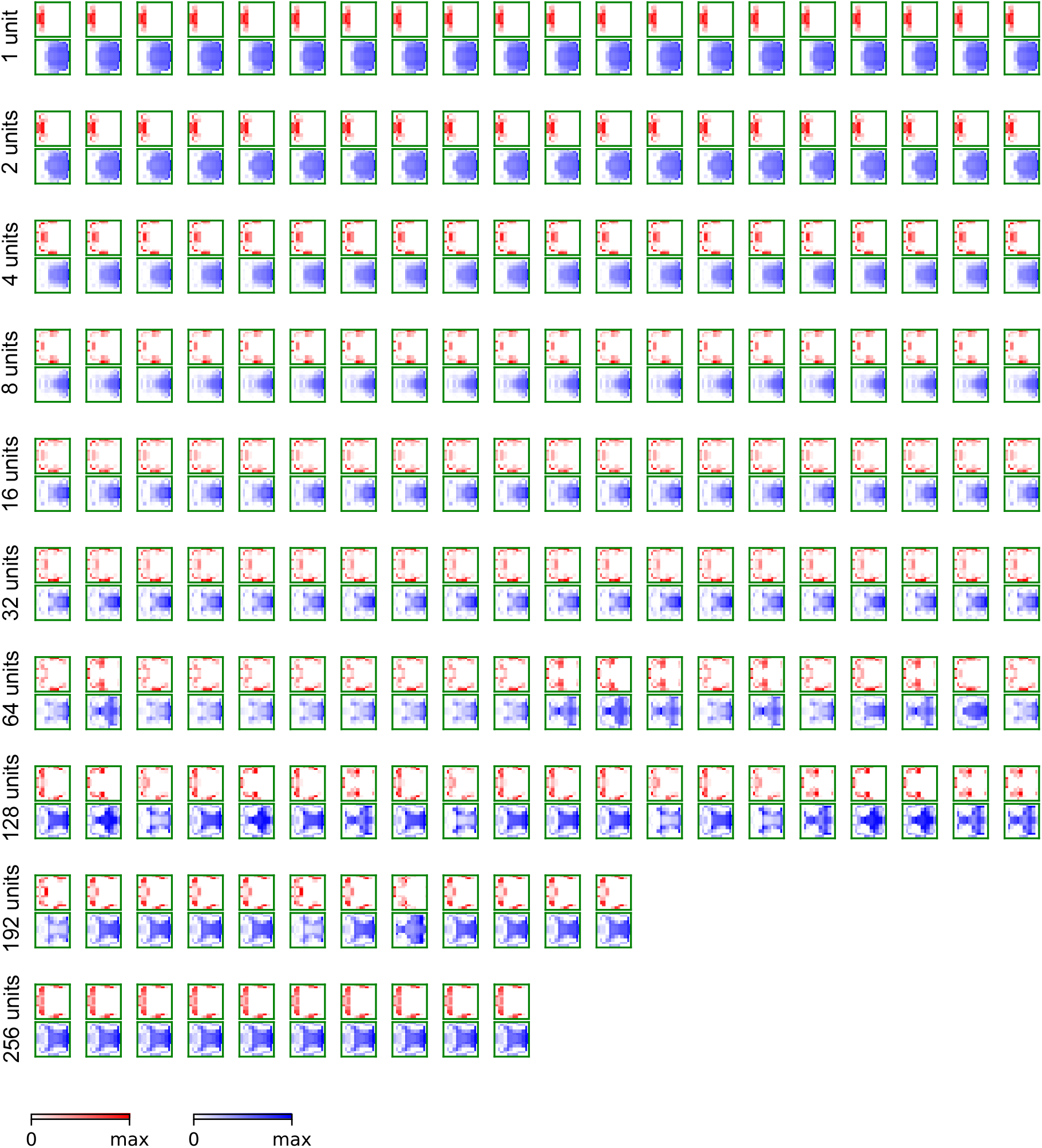
Inward solutions for models with different numbers of units.

**Figure 8–Figure supplement 1.**
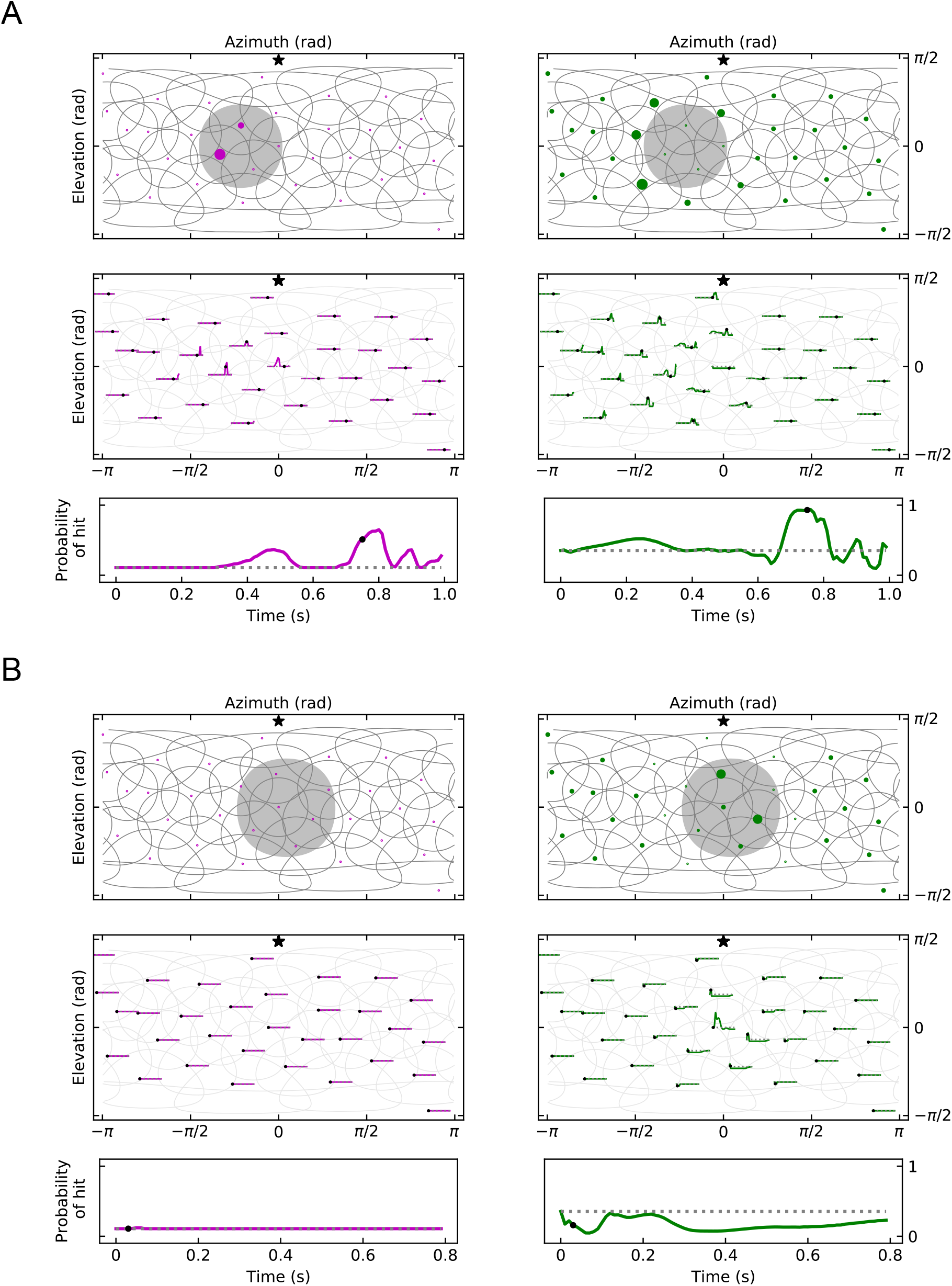
(A) An example of response patterns to a miss stimulus (Video 7 and Video 8). (B) An example of response patterns to a retreat stimulus (Video 9 and Video 10). The snapshots of the rotational case are not shown, but can be found in the videos (Video 11 and Video 12).

**Figure 8–Figure supplement 2.**
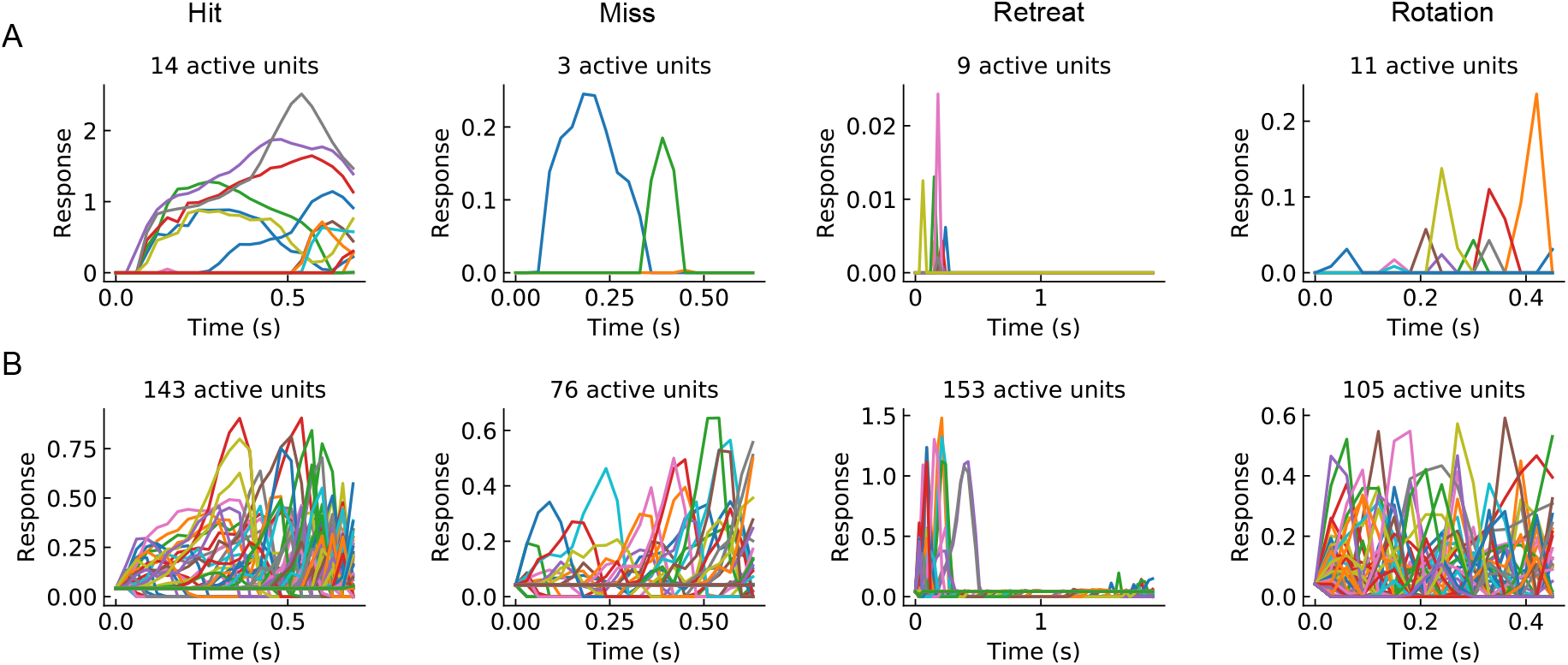
(A) Sample response curves of the active units in the outward solution with *M* = 256 for different types of stimuli (from left to right: hit, miss, retreat, and rotation) (B) As in (A), but for an inward solution.

**Figure 10–Figure supplement 1.**
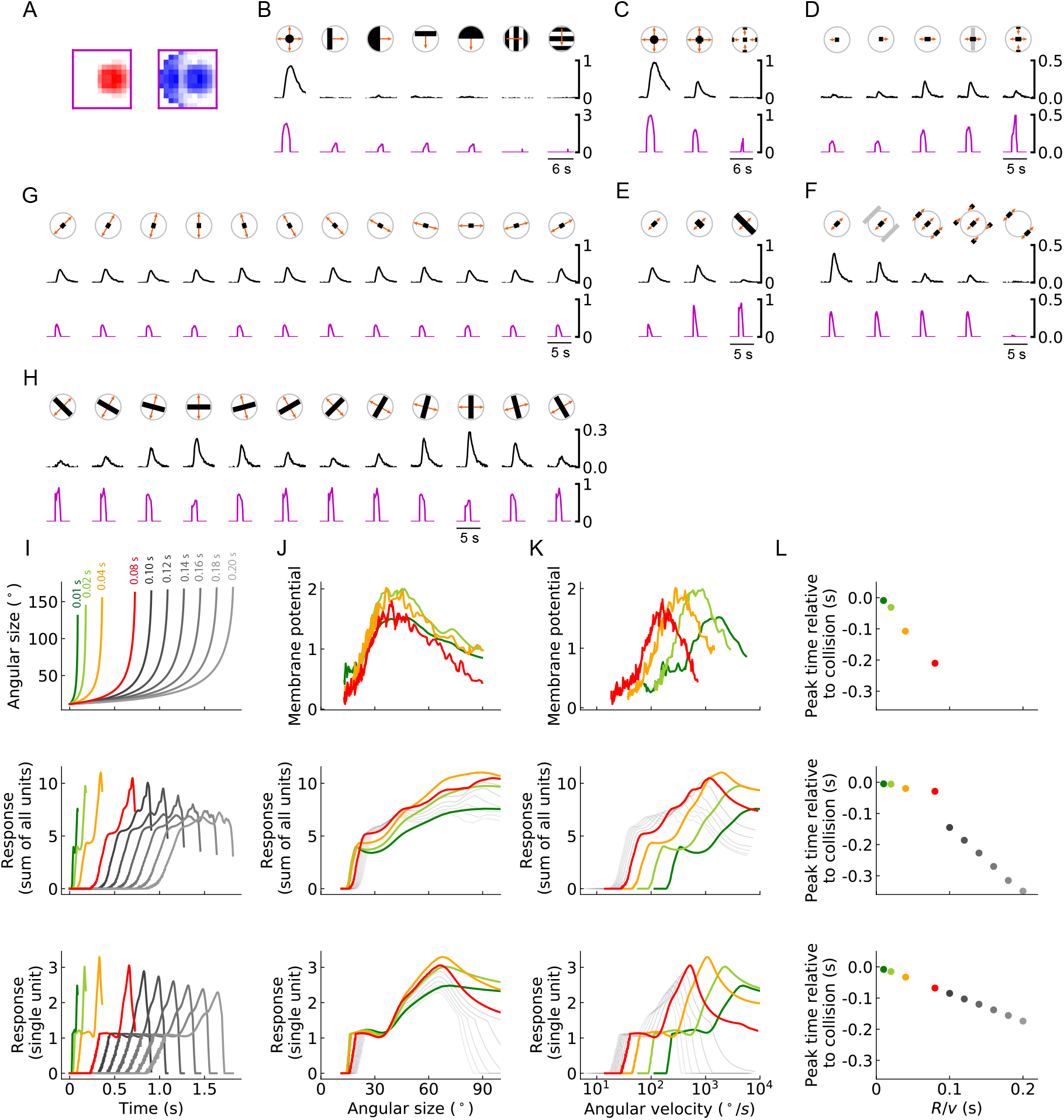
The same as the main figure for a different trained outward model, the filters of which are shown in (A).

